# Notch-Jagged signaling complex defined by an interaction mosaic

**DOI:** 10.1101/2021.02.19.432005

**Authors:** Matthieu R. Zeronian, Oleg Klykov, Júlia Portell i de Montserrat, Maria J. Konijnenberg, Anamika Gaur, Richard A. Scheltema, Bert J.C. Janssen

**Affiliations:** Crystal and Structural Chemistry, Bijvoet Center for Biomolecular Research, Department of Chemistry, Faculty of Science, Utrecht University, Padualaan 8, 3584 CH Utrecht, The Netherlands; Biomolecular Mass Spectrometry and Proteomics, Bijvoet Center for Biomolecular Research and Utrecht Institute for Pharmaceutical Sciences, Utrecht University, Padualaan 8, 3584 CH Utrecht, The Netherlands.; Netherlands Proteomics Centre, Padualaan 8, 3584 CH Utrecht, The Netherlands.

**Keywords:** Notch1, Jagged1, cell signaling, EGF repeats, protein-protein interaction, glycoprotein, cross-linking mass spectrometry, XlinkX/PD, PhoX

## Abstract

The Notch signaling system links cellular fate to that of its neighbors, driving proliferation, apoptosis, and cell differentiation in metazoans, whereas dysfunction leads to debilitating developmental disorders and cancers. Other than a five-by-five domain complex, it is unclear how the 40 extracellular domains of the Notch1 receptor collectively engage the 19 domains of its canonical ligand Jagged1 to activate Notch1 signaling. Here, using cross-linking mass spectrometry (XL-MS), biophysical and structural techniques on the full extracellular complex and targeted sites,we identify five distinct regions, two on Notch1 and three on Jagged1, that form an interaction network.The Notch1 membrane-proximal regulatory region individually binds to the established Notch1 epidermal growth factor (EGF) 8-13 and Jagged1 C2-EGF3 activation sites, as well as to two additional Jagged1 regions, EGF 8-11 and cysteine-rich domain (CRD). XL-MS and quantitative interaction experiments show that the three Notch1 binding sites on Jagged1 also engage intramolecularly.These interactions, together with Notch1 and Jagged1 ectodomain dimensions and flexibility determined by small-angle X-ray scattering (SAXS), support the formation of backfolded architectures. Combined, the data suggest that critical Notch1 and Jagged1 regions are not distal, but engage directly to control Notch1 signaling, thereby redefining the Notch1-Jagged1 activation mechanism and indicating new routes for therapeutic applications.

---

Notch signaling plays a central role in developmental processes by determining cell fate decisions in tissues during development. In adults, these signals both determine differentiation and maintenance of neuronal and hematopoietic stem cells as well as regulate the immune system (Ables *et al*., 2011; Bray, 2016; Louvi and Artavanis-Tsakonas, 2006; Radtke *et al*., 2013). Dysregulation often leads to debilitating diseases in humans, including congenital disorders and cancers (Aster *et al*., 2017; Mašek and Andersson, 2017; Siebel and Lendahl, 2017; Weng *et al*., 2004). The mammalian Notch1 receptor is the prototypical member of the Notch protein family, which consists of four paralogs (Notch1-4) that all receive signals from the associated ligands Jagged1, Jagged2, Delta-like1, and Delta-like4: in trans (from adjacent cells) to initiate signaling, or in cis (from the same cell) to inhibit signaling. The Notch1-Jagged1 receptor-ligand pair has been widely studied at functional, cellular, and molecular levels (Bray, 2016; Siebel and Lendahl, 2017). Both Notch1 and Jagged1 are type-I transmembrane proteins with large modular extra-cellular segments that determine interaction specificity and control the activation of signaling. Notch1 has an extracellular segment of 209 kDa composed of 36 EGF repeats followed by the NRR at the membrane-proximal side, and differs from its paralogs in the number of EGF domains: from 36 for Notch2, 34 for Notch3 and 29 for Notch4. The Jagged1 ectodomain (139 kDa) is similar to that of Jagged2 and is composed of a C2 lipid-binding domain, a Delta/Serrate/Lag-2 (DSL) domain, 16 EGF repeats and a CRD at the membrane-proximal side.

The prevailing model for canonical Notch activation states that ligand binding at Notch1 EGF8-12 and an endocytosis-induced pulling force (Chowdhury *et al*., 2016; Gordon *et al*., 2015; Lovendahl *et al*., 2018; Luca *et al*., 2017; Meloty-Kapella *et al*., 2012; Rebay *et al*., 1991; Seo *et al*., 2016; Wang and Ha, 2013), generated by the signal-sending cell on the Notch-ligand complex (Nichols *et al*., 2007; Parks *et al*., 2000), triggers a conformational change and proteolytic processing in the Notch NRR located 24 EGF domains downstream of the ligand binding site (Brou *et al*., 2000; Gordon *et al*., 2007; Mumm *et al*., 2000). After Notch cleavage within the transmembrane domain (Kopan *et al*., 2009; Yang *et al*, 2019), the Notch intracellular domain translocates to the nucleus where it regulates transcription (Bray et al, *et al*, 2018). At the N-terminal side of Jagged1, the C2-EGF3 region is important for Notch1 binding (Chillakuri *et al*., 2013; Cordle *et al*., 2008a; Luca *et al*., 2017; Shimizu *et al*., 1999; Suckling *et al*., 2017). A recent structural study demonstrated that the Notch1 EGF8-12 region interacts in an antiparallel fashion through an extended interface with the Jagged1 C2-EGF3 region (Luca *et al*., 2017). Additional interactions add complexity to the mechanism of Notch activation and regulation. Notch-ligand, Notch-Notch and ligand-ligand interactions in cis can both inhibit (Del Á lamo *et al*., 2011; D’Souza *et al*., 2008; Sprinzak *et al*., 2010) or activate (Nandagopal *et al*., 2019) signaling. In addition to the canonical ligand binding site on EGF8-12 and the conformational change in the NRR, several other extracellular regions, such as EGF6, EGF25-26 and EGF36, seem to play a role in Notch function (Holdener and Haltiwanger, 2019; Kakuda and Haltiwanger, 2017; Lawrence *et al*., 2000; Pei and Baker, 2008; Rampal *et al*., 2005; Sharma *et al*., 2013; Xu *et al*., 2005). Also, the Jagged1 extracellular segment harbors additional functionality other than the C2-EGF3 region interacting to Notch. It has been suggested that Jagged and Delta-like C2 domain binding to membranes has an important role in regulating ligand-dependent Notch signaling (Chillakuri *et al*., 2013; Suckling *et al*., 2017). The CRD in Xenopus Serrate-1, a homolog of mammalian Jagged1, is required for Notch activation in primary neurogenesis (Kiyota and Kinoshita, 2002). These studies indicate that several sites in the Notch and Jagged extracellular segments may contribute to Notch signaling and regulation.

Structural studies have revealed details of key interaction sites (Luca *et al*., 2015, 2017) and indicate that flexibility is present to a certain extent in the Notch and Jagged ectodomains (Suckling *et al*., 2017; Weisshuhn *et al*., 2016). A low-resolution negative stain electron microscopy reconstruction of the Notch1 ectodomain resolved distinct globular dimer states, although this protein was purified in an unconventional manner (Kelly *et al*., 2010). Backfolded models for the Notch ectodomain have also been suggested based on genetic and interaction studies (Pei and Baker, 2008; Sharma *et al*., 2013; Xu *et al*., 2005). Nonetheless, direct observations of ectodomain flexibility and backfolding are limited. While Notch-Jagged interaction studies have focused predominantly on the well-established Notch1 EGF11-12 - Jagged1 C2-EGF3 regions, other sites may play a direct role in this intermolecular interaction. Structural and biophysical studies on the full extracellular portions of Notch and Jagged have however been limited due to the size, flexibility and low expression levels of the proteins, hampering the identification of several interacting regions.

In this study, we combine cross-linking mass spectrometry, quantitative interaction assays and small-angle X-ray scattering (SAXS) on purified Notch1 and Jagged1 full ectodomains, as well as shorter constructs, to probe the structure of the Notch1-Jagged1 complex and of the unliganded proteins (Fig. 1a-d). This analysis reveals several, hitherto unreported, intra- and intermolecular interaction regions. We show that Jagged1 C2-EGF3, EGF8-11 and CRD can all interact with Notch1 EGF33-NRR and that the Notch1 NRR is sufficient for the interaction with Jagged1 C2-EGF3. In addition, the Notch1 EGF8-13 region directly interacts with Notch1 EGF33-NRR. XL-MS analysis shows that four regions, C2-EGF1, EGF5-6, EGF9-12 and CRD, are in proximity within Jagged1, and we confirmed direct interactions for C2-EGF3 binding to EGF8-11 and to CRD. These data, together with SAXS analysis of the Notch1 and Jagged1 ectodomains, suggest that the proteins are backfolded and indicate that regions in both proteins, *i*.*e*. Notch1 EGF8-13, Notch1 EGF33-NRR and Jagged1 C2-EGF3, previously shown to be important for Notch signaling, affect each other directly.

**Figure 1.**
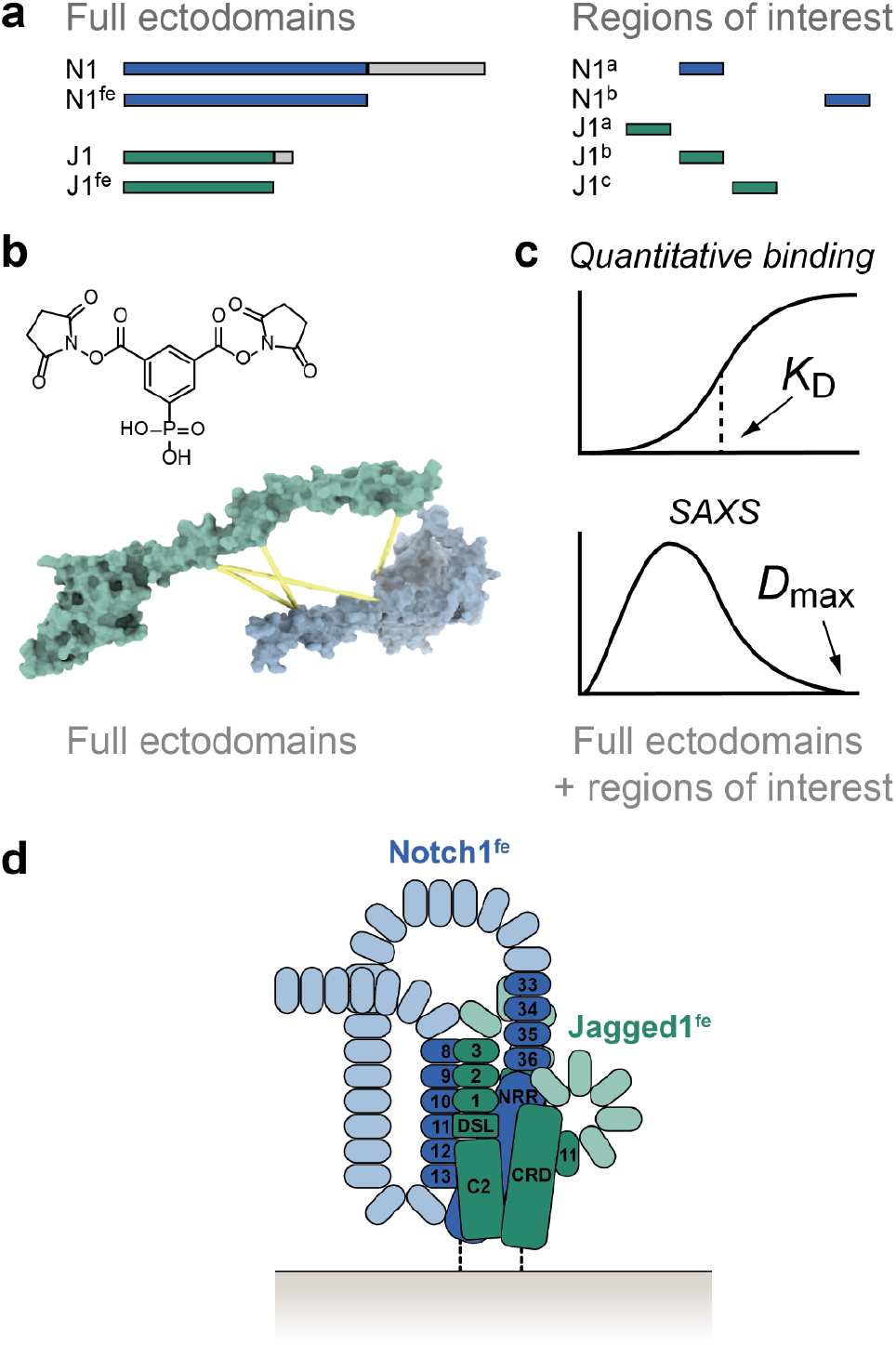
XL-MS and biophysical studies reveal an interaction network in the Notch1-Jagged1 complex. **a** Notch1^fe^, Jagged1^fe^ and targeted sites are expressed in HEK293 cells and purified by IMAC and SEC. **b** Identification of regions in proximity in the Notch1^fe^-Jagged1^fe^ complex by XL-MS using PhoX (Steigenberger *et al*., 2019). **c** The purified full ectodomain samples and shorter regions of interest are used in quantitative interaction experiments and SAXS studies. **d** The resulting data provides insights into the molecular architecture of the Notch1-Jagged1 complex, represented here in a *cis* setting.

## Results

### XL-MS of the Notch1-Jagged1 complex reveals a mosaic of interaction sites

To determine which regions, beyond the canonical Notch1^EGF8-12^-Jagged1^C2-EGF3^ interaction site, are involved in receptor-ligand binding, we probed full ectodomains of Notch1 and Jagged1 (Notch1^fe^-Jagged1^fe^) with XL-MS (Figures 1, 2a, 2b and Tables S1-S3). Two variants of Jagged1 were used: a wild-type version (Jagged1^fe,*wt*^), and one with five point mutations in the Jagged C2 region (Jagged1^fe,HA^) that provide higher-affinity binding to Notch1 EGF8-12 when incorporated in a Jagged1 C2-EGF3 construct (Luca *et al*., 2017). In surface plasmon resonance (SPR) experiments, where Notch1^fe^ is coupled at the C-terminus to the sensor surface to achieve a close-to-native topology (see Methods), Notch1^fe^-Jagged1^fe,HA^ interact with a dissociation constant (*K*_D_) of 1 µM and Jagged1^fe,HA^ interacts with similar affinity to the EGF8-13 portion of Notch1, while no interaction was measured between Jagged1^fe,*wt*^ and Notch1 EGF8-13 at 1 µM (Figures 2c, 2d, S1a and S1b).

**Figure 2.**
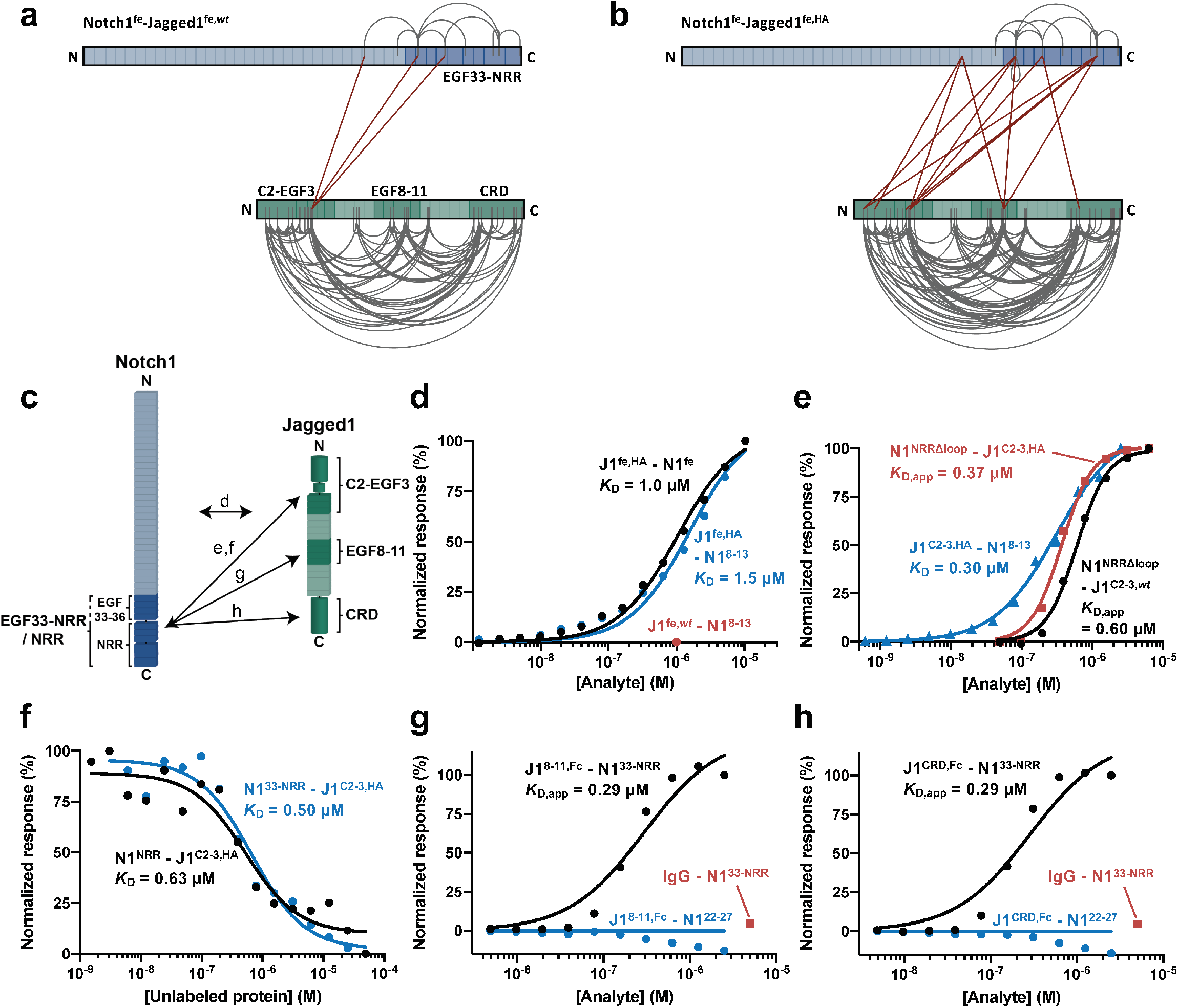
The Notch1 C-terminal region interacts with Jagged1^C2-EGF3^, Jagged1^EGF8-11^ and Jagged1^CRD^ in the Notch1^fe^-Jagged1^fe^ complex. **a-b** Overview of the detected distance constraints from the XL-MS experiments, for wild-type **a** and high-affinity **b** versions of Jagged1^fe^. **c** Schematic representation of the interactions reported in panels **d-h**, based on the XL-MS data and quantitative binding experiments. **d** SPR equilibrium binding plots of Jagged1^fe,HA^ to Notch1^fe^ (black) and to Notch1^EGF8-13^ (blue). Jagged1^fe,*wt*^ does not interact with Notch1^EGF8-13^ at 1 µM (red). **e** SPR equilibrium binding plots of Notch1^NRRΔloop^ to wild-type (black) and to high-affinity (red) versions of Jagged1^C2-EGF3^, and of Jagged1^C2-EGF3,HA^ to Notch1^EGF8-13^ (blue). A Hill coefficient of 2 is used to model the Notch1^NRRΔloop^ - Jagged1^C2-EGF3^ interactions (see also Methods). **f** MST binding curve of Notch1^NRR^ (black) and Notch1^EGF33-NRR^ (blue) to Jagged1^C2-EGF3,HA^. **g-h** SPR equilibrium binding plots indicate interaction of dimerized Jagged1^EGF8-11,Fc^ **g** and dimerized Jagged1^CRD,Fc^ **h** to Notch1^EGF33-NRR^ (black) but not to Notch1^EGF22-27^ that acts as negative control (blue). The Fc domain does not interact with Notch1^EGF33-NRR^ as shown by the IgG control at 5 µM (red). See also Figures S1-S5 and Table 1.

Purified Notch1 and Jagged1 full ectodomain proteins were incubated at a 1 to 1 molar ratio to induce complex formation, *i*.*e*. Notch1^fe^-Jagged1^fe,*wt*^ and Notch1^fe^-Jagged1^fe,HA^, and cross-linked with the lysine-targeting PhoX cross-linking reagent (Steigenberger *et al*., 2019). In subsequent steps, the samples were subjected to deglycosylation, enriched for cross-linked peptides by immobilized metal affinity chromatography (IMAC) and finally analyzed by liquid chromatography coupled to tandem mass spectrometry (LC-MS/MS). From three independent replicates for each complex, we detected 166 unique distance restraints for Notch1^fe^-Jagged1^fe,*wt*^ and 232 for Notch1^fe^-Jagged1^fe,HA^. As an additional step to reduce false positives and remove distance constraints arising from non-specific aggregation we solely retained restraints detected in at least two out of three replicates (Klykov *et al*., 2020). This reduced the output to 113 and 164 restraints for Notch1^fe^-Jagged1^fe,*wt*^ and Notch1^fe^-Jagged1^fe,HA^ respectively (Figures 2a and 2b). For both complex samples, few intra-links were detected for Notch1^fe^ (9 for Notch1^fe^-Jagged1^fe,*wt*^ and 12 for Notch1^fe^-Jagged1^fe,HA^, Figures S2a and S2d). The number of intra-links for Jagged1^fe^ was however significantly larger and increased by 38% for the mutant (100 for Notch1^fe^-Jagged1^fe,*wt*^ and 138 for Notch1^fe^-Jagged1^fe,HA^). A similar trend was visible in the number of intermolecular connections between Notch1 and Jagged1 where 3 inter-links were detected for Notch1^fe^-Jagged1^fe,*wt*^ and 13 for Notch1^fe^-Jagged1^fe,HA^. This identification of intra- and inter-links suggests that the mutant protein, Jagged1^fe,HA^, assisted by the stronger interaction between the two molecules, adopts a less flexible conformation compared to Jagged1^fe,*wt*^, and provides more efficient complex formation that is beneficial for our approach (Fürsch *et al*., 2020).

The inter-links reveal that in the Notch1^fe^-Jagged1^fe^ complex, three Jagged1 regions, C2-EGF1, EGF10 and CRD are in proximity to the Notch1 EGF29-NRR site with most inter-links arising from the Jagged C2-EGF1 region. The XL-MS experiments do not reveal any cross-links or mono-links between Notch1 EGF8-12 and Jagged1 C2-EGF3 (Figures 2a, 2b and S2a), the well-established interaction site (Luca *et al*., 2017) for which we find a *K*_D_ of 0.3 µM by SPR, using the high-affinity variant of Jagged1 C2-EGF3 (Figures 2e and S1c). There are two possible explanations for the lack of links to Notch1 EGF8-12. (I) The two lysine residues in Notch1 EGF8-12, Lys395 and Lys428, are occluded in the Notch1^fe^-Jagged1^fe^ complex or (II) the lysines are occluded from the cross-linking reaction by O-linked glycans. Shotgun mass spectrometric analysis of non-cross-linked Notch1^fe^ covers the segment containing the two lysines within the Notch1 EGF8-12 region, indicating that the relevant peptides can be identified (Figure S2a). A large part of the Notch1^fe^ EGF repeat region is decorated with O-linked glycosylation sites, with an average of 1.5 sites per EGF domain based on sequence prediction (Takeuchi and Haltiwanger, 2014), and we cannot fully exclude the glycans prevent the cross-linking reaction. Notably, however, 25 cross-links are identified in the Notch1 EGF29-36 region, predicted to contain slightly less O-linked glycosylation sites, *i*.*e*. 1.1 sites per EGF domain (Takeuchi and Haltiwanger, 2014). Combined, these observations suggest that Notch1 EGF8-12 is hidden in the folded Notch1 full ectodomain. Although the XL-MS analysis has not revealed all the interacting regions on Notch1 in the Notch1^fe^-Jagged1^fe^ complex, it does indicate that the Notch1 C-terminal region plays an important role in the interaction with Jagged1.

### Notch1 NRR directly interacts with Jagged1 C2-EGF3

To further investigate interacting regions, we generated shorter Notch1 and Jagged1 constructs (Figure S3) and probed them by SPR and microscale thermophoresis (MST). The Notch1^EGF33-NRR^ site interacts directly with Jagged1^C2-EGF3^ in MST (Figures 2f and S1g) and in SPR (Figures S4a-S4c), and this interaction is independent of the high-affinity mutations in the C2 domain of Jagged1 (Table 1; Figures S4a-S4d). Jagged1^C2-EGF3^ is required and sufficient for the interaction with Notch^EGF33-NRR^ (Figures S4a-S4f). The Notch1^EGF33-NRR^-Jagged1^C2-EGF3^ binding site was further defined to Notch1^NRR^, that binds with a *K*_D_ of 0.6 µM to Jagged1^C2-EGF3,HA^, measured in solution by MST (Figures 2f and S1f). In the NRR, a large unstructured loop (consisting of 38 residues) that contains the heterodimerization S1 cleavage site (Gordon *et al*., 2007; Logeat *et al*., 1998) is not required for interaction (Figures 2e and S1d). In addition, the interaction is not affected by the high-affinity mutations in Jagged1^C2-EGF3^, as the *K*_D_ values determined by SPR for Notch1^NRRΔloop^ binding to Jagged1^C2-EGF3,*wt*^ or to Jagged1^C2-EGF3,HA^ are similar (Figures 2e, S1d and S1e). Docking of the Notch1^NRR^-Jagged1^C2-EGF3^ complex, using the intermolecular cross-links as restraints, suggests that the Notch1 NRR engages domains DSL and EGF1 of Jagged1 (Figure S5). Taken together, our interaction data on the smaller Notch1 and Jagged1 portions show that the Notch1 NRR is responsible for the interaction with the Jagged1 C2-EGF3 region.

**Table 1.**
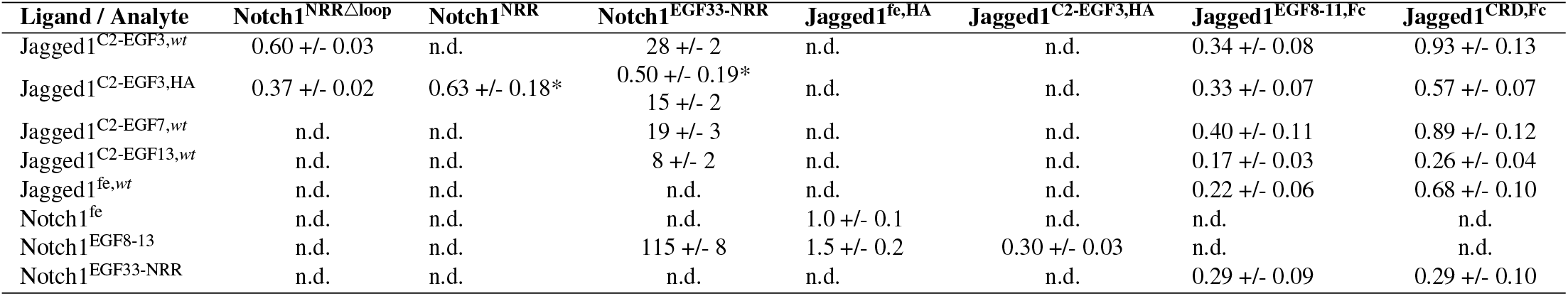
Summary of measured affinities. All values are expressed in µM and derived from MST (indicated by an asterisk) or SPR experiments. n.d. = not determined. Related to Figures 2-4, S1, S3 and S4.

### Notch1^EGF33-NRR^ contains low affinity sites for Jagged1^EGF8-11^ and Jagged1^CRD^

The XL-MS data of Notch1^fe^-Jagged1^fe,HA^ indicates that two additional regions in Jagged1, EGF10 and CRD, are in proximity to the Notch1 EGF33-NRR site (Figure 2b). SPR binding experiments confirm the direct interactions to Notch1^EGF33-NRR^, albeit with much lower affinity than the Jagged1 C2-EGF3 region, with no binding of Jagged1^EGF8-11^ or Jagged1^CRD^ to Notch1^EGF33-NRR^ observed at concentration of 20 µM (data not shown). To enhance a possible weak affinity, we employed a widely used strategy for cell and surface binding assays of artificially dimerizing proteins (Czajkowsky *et al*., 2012) that has previously been used to measure Notch interactions (Sharma *et al*., 2013; Shimizu *et al*., 1999). Fc-tagged versions of Jagged1^EGF8-11^ and Jagged1^CRD^, that are covalently dimerized by the Fc tag, interact both with a *K*_D,app_ of 0.29 µM to Notch1^EGF33-NRR^ (Figures 2g, 2h, S1h and S1i).

### Notch1^fe^ is flexible and has intramolecular interactions

SAXS analysis coupled to size-exclusion chromatography (SEC-SAXS) shows that monomeric Notch1^fe^ is a flexible molecule (Figures 3a and 3b), has a radius of gyration (*R* _g_) of 105 +/− 0.4 Å (Figure 3c) and a maximum distance (*D*_max_) of 380 Å (Figure 3d). This suggests that Notch1^fe^ does not exist as an elongated molecule, as it would have a *D*_max_ of 1,027 Å for a fully elongated Notch1^fe^ (see Methods), but instead has considerable backfolding. Backfolded models were previously suggested based on genetic (Xu *et al*., 2005) and interaction data (Pei and Baker, 2008; Sharma *et al*., 2013), where the EGF domain connections were determined to confer flexibility to the Notch1 extracellular region (Weisshuhn et al., 2016). In addition, two parts in Notch1, the EGF domain connections were determined to confer flexibility to the Notch1 extracellular region (Weisshuhn et al., 2016). In addition, two parts in Notch1, EGF8-13 and EGF33-NRR, interact with a *K*_D_ of 115 μM (Figures 3e and S1j). While this is a relatively low affinity for an intermolecular interaction, *i.e.* as in a Notch1 dimer, it may be possible that these regions interact directly in an intramolecular fashion within the same Notch1 molecule. Overall, the backfolding suggests that EGF domains may become buried in the fully folded molecule, providing further support to the data obtained by XL-MS.

**Figure 3.**
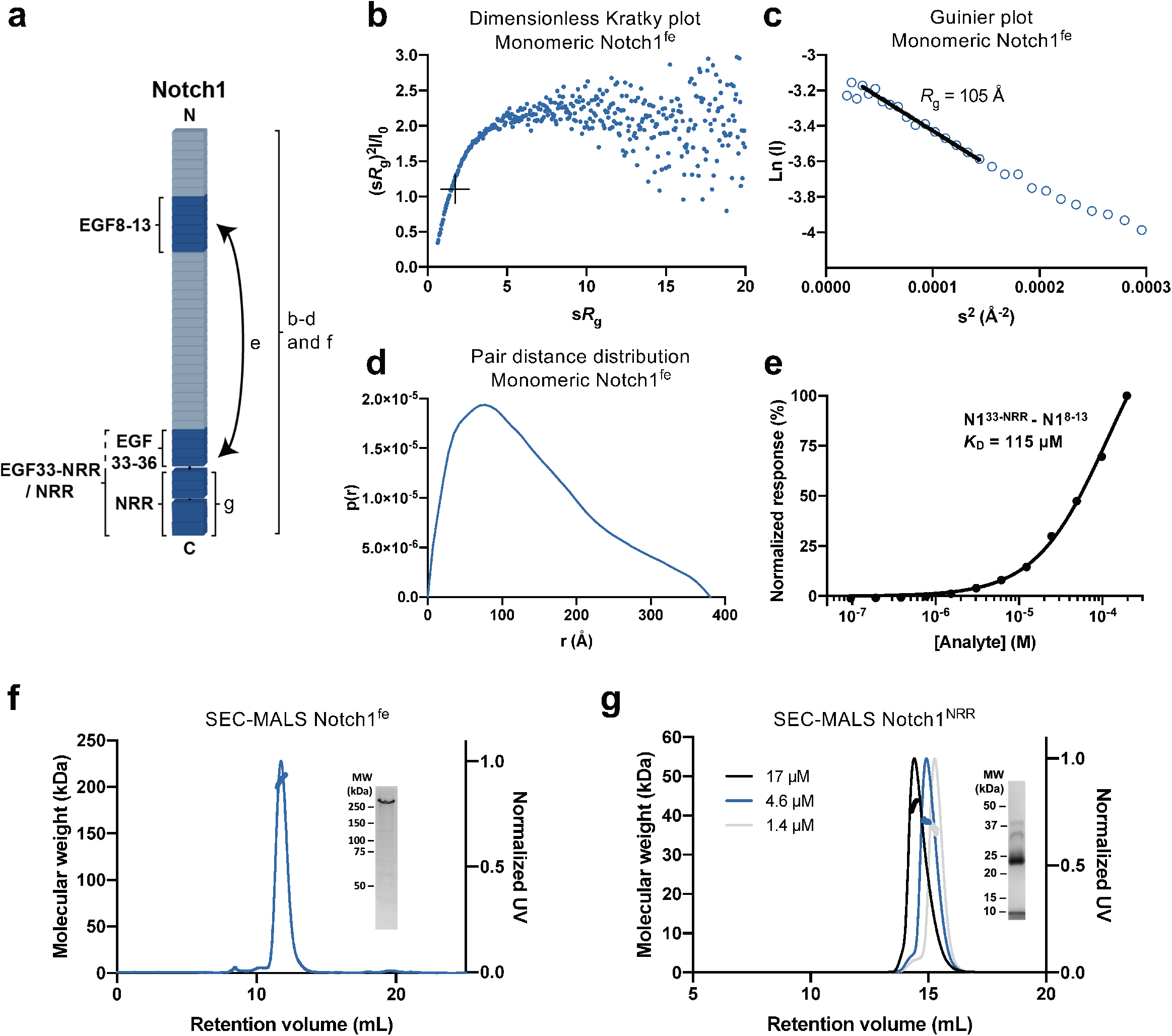
Notch1^fe^ is flexible and the NRR dimerizes weakly. **a** Schematic representation of the interaction and biophysical experiments on regions reported in panels **b-g. b-d** Structural analysis of monomeric Notch1^fe^ from SEC-SAXS, including Dimensionless Kratky plot with crosshairs indicating the peak position for a globular protein **b**, Guinier plot with a black line indicating the fit used to derive the *R*_g_ **c** and pair distance distribution function **d. e** SPR equilibrium binding plot of Notch1^EGF33-NRR^ to Notch1^EGF8-13^. **f** SEC-MALS analysis of Notch1^fe^ shows a monomeric and monodisperse sample (thick lines indicate the molecular weight, left axis). Inset: Coomassie-stained SDS-PAGE of purified Notch1^fe^ in reducing conditions. **g** SEC-MALS analysis of Notch1^NRR^ at three concentrations determined at elution shows a monomer-dimer equilibrium (thick lines indicate the molecular weight, left axis). Inset: Coomassie-stained SDS-PAGE of purified Notch1^NRR^ in reducing conditions, note that Notch1^NRR^ is processed at the S1 cleavage site into two fragments of 8 kDa and 27 kDa. See also Figures S1, S3, S6 and Tables 1 and 2.

### Notch1 dimerizes through the NRR

Notch1^fe^ is a monomer at a concentration of 0.26 µM and has a molecular weight of 209 +/− 2.4 kDa (Figure 3f). This correlates well with the theoretical molecular weight of 200-220 kDa that is dependent on the glycosylation state (Kakuda and Haltiwanger, 2017; Taylor *et al*., 2014). Interestingly, our XL-MS data showed that Notch1^fe^ can form dimers, which can be detected by XL-MS when the same residue in the protein sequence is linked by two different peptides induced by *e*.*g*. a missed cleavage. One self-link at lysine residue 1314 in EGF34 arises from an intermolecular Notch1-Notch1 interaction (Figure 2b). In addition, the Notch1 NRR itself (Notch1^NRR^) undergoes weak concentration-dependent dimerization during size-exclusion chromatography coupled to multi-angle light scattering (SEC-MALS) analysis at concentrations ranging from 1.4 to 17 µM (Figure 3g). Dimerization of the NRR has previously been reported for Notch3 and was predicted for the Notch1 NRR based on similarities in crystal packing comparing the NRR of Notch3 and Notch1 (Gordon *et al*., 2009b, 2009a; Xu *et al*., 2015). The NRR-controlled dimerization of Notch3 may maintain the receptor in an autoinhibited state before ligand binding (Xu *et al*., 2015). We determined a crystal structure of the S1-cleaved mouse Notch1 NRR (Figures S6a-S6c; PDB: 7ABV) that shows the same dimerization interface as its human ortholog (Gordon *et al*., 2009b, 2009a). N-linked glycans, that do not seem to interfere with dimerization, are visible in the electron density at position N1489, as also reported previously (Wu *et al*., 2010), and additionally at position N1587 (Figure S6a). Taken together, the XL-MS analysis on Notch1^fe^ and dimerization of Notch1^NRR^ indicate that Notch1 can dimerize through the membrane proximal region.

### Jagged1^fe^ is backfolded and oligomerizes

Jagged1^fe,HA^ has a weak propensity to dimerize. Up to a concentration of 1.6 µM, Jagged1^fe,HA^ is a monomer with a molecular weight of 137 +/− 0.2 kDa (Figures 4a and 4b) that correlates well with the theoretical molecular weight of 120-140 kDa depending on the glycosylation state (Pandey *et al*., 2020; Thakurdas *et al*., 2016). At higher concentrations, Jagged1^fe^ forms oligomers (Figures 4c-4e). In sedimentation velocity analytical ultracentrifugation (SV-AUC), at 5 µM 19 % of Jagged1^fe,HA^ consists of oligomers, and this increases to 31 % at 20 µM (Figure 4c). Concentration-dependent dimerization is also supported by batch SAXS analysis. At 5 µM the *R*_g_ of Jagged1^fe,HA^ is 81.2 +/− 0.8 Å and this increases to 102 +/− 0.4 Å at 42 µM (Table 2; Figure 4d) indicating more Jagged1^fe,HA^ dimers or larger oligomeric species are present at higher concentration.

**Figure 4.**
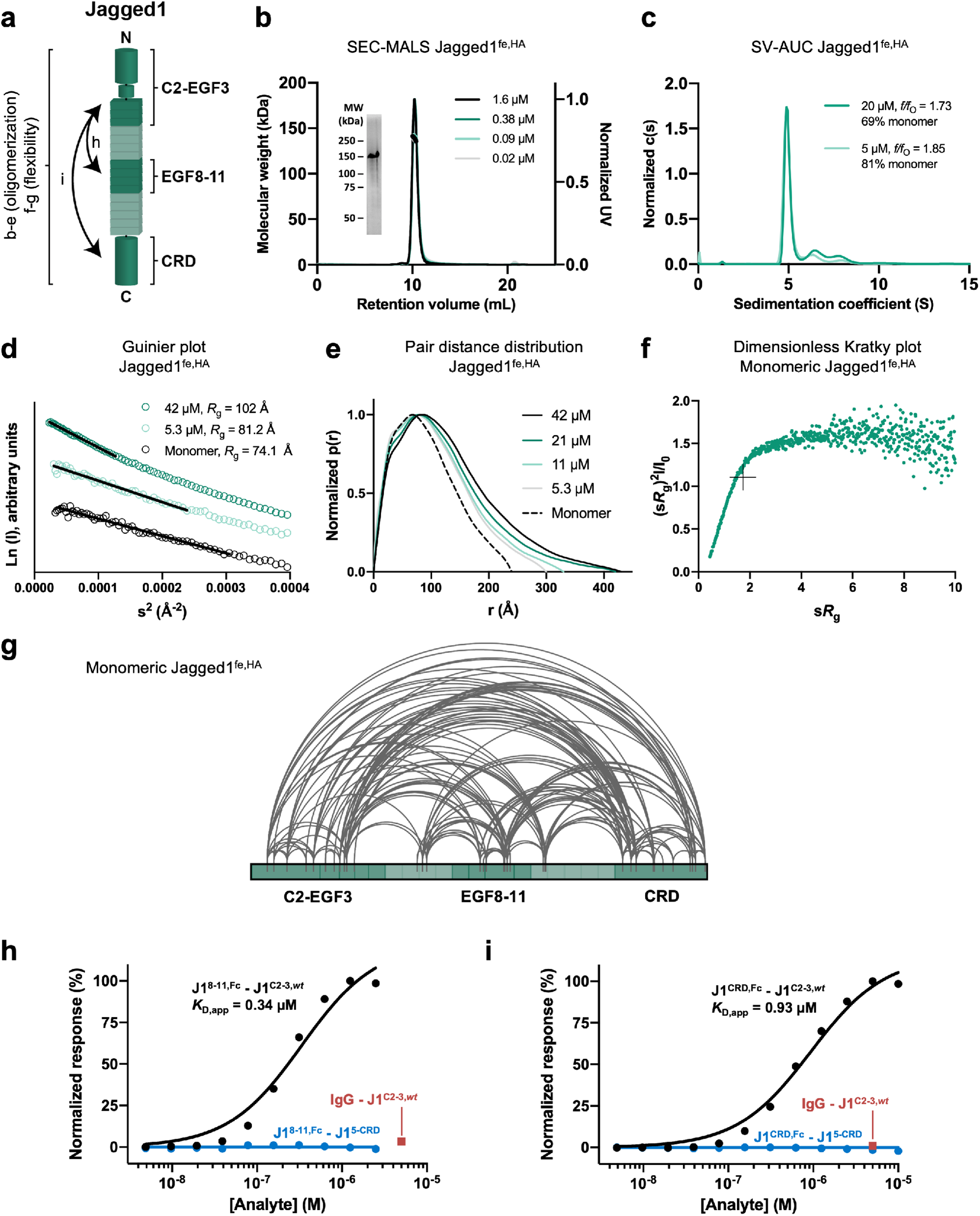
Jagged1^fe,HA^ is backfolded, flexible and oligomerizes weakly. **a** Schematic representation of the interactions and biophysical experiments on regions reported in panels **b-j. b** SEC-MALS analysis of Jagged1^fe,HA^ at four concentrations determined at elution shows overlapping monomeric and monodisperse peaks (thick lines indicate the molecular weight, left axis). Inset: Coomassie-stained SDS-PAGE of the purified sample in reducing conditions. **c** SV-AUC analysis shows that Jagged1^fe,HA^ oligomerizes in a concentration-dependent manner. **(d-f)** SAXS analysis of Jagged1^fe,HA^ in batch and from monomeric SEC-SAXS fractions including Guinier plot with black lines indicating the fits used to derive the *R*_g_ **d**, pair distance distribution function **e** and dimensionless Kratky plot with crosshairs indicating the peak position for a globular protein **f. g** Overview of the detected distance constraints from the XL-MS experiments for monomeric Jagged1^fe,HA^. **h-i** SPR equilibrium binding plots indicate interaction of Jagged1^EGF8-11,Fc^ **h** and Jagged1^CRD,Fc^ **i** to Jagged1^C2-EGF3^ (black) but not to Jagged1^EGF5-CRD^ that acts as negative control (blue). The Fc domain does not interact with Jagged1^C2-EGF3,*wt*^ as shown by the IgG control at 5 µM (red). See also Figures S1, S2, S3, S7 and Tables 1 and 2.

**Table 2.**
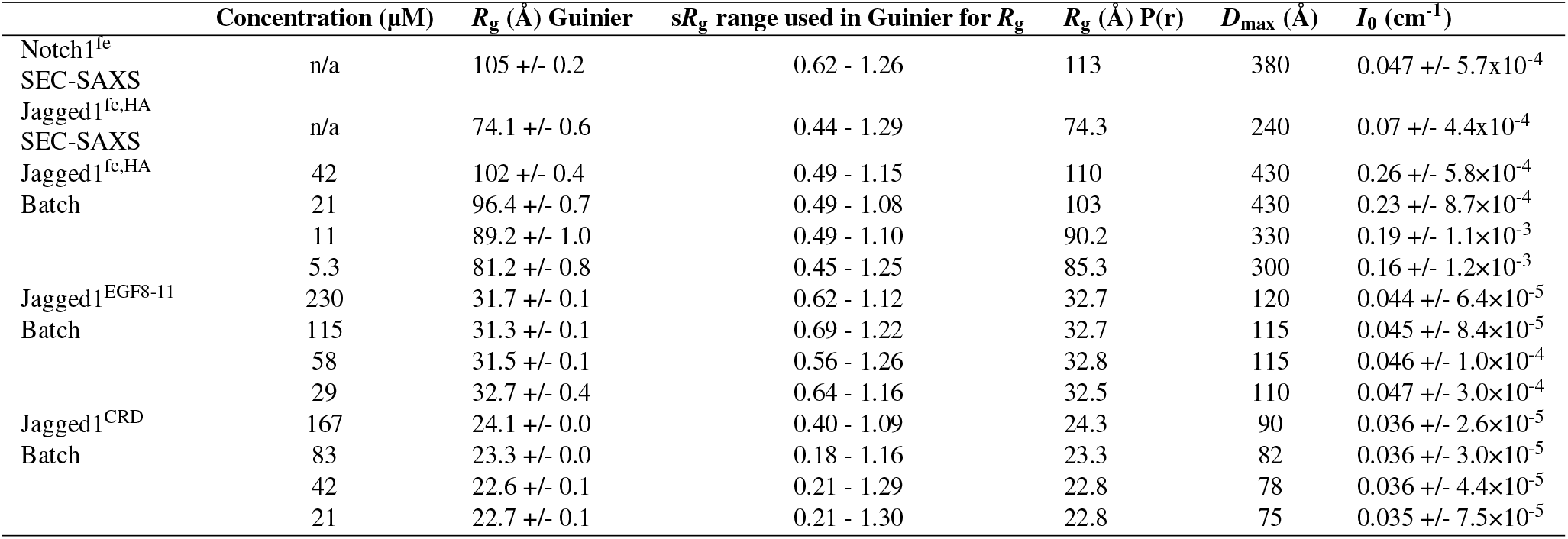
Structural parameters derived from SAXS experiments. SAXS batch data I0 have been normalized by the sample concentration to allow for comparison between samples. Non-normalized *I*_0_ values are available on SASBDB under the accession codes defined in Data and materials availability. n/a = not applicable. Related to Figures 3, 4 and S7.

We used SEC-SAXS to separate monomeric Jagged1^fe,HA^ from oligomeric species. The region at the right side of the Jagged1^fe,HA^ elution peak, *i*.*e*. at larger retention volume, was selected for further analysis as this region most likely represents a monomeric fraction. Jagged1^fe,HA^ has a *R*_g_ of 74.1 +/− 0.6 Å (Figure 4d) and a *D*_max_ of 240 Å (Figure 4e). The normalized Kratky plot indicates that structural flexibility is present in the Jagged1 ectodomain (Figure 4f). SAXS analysis of smaller Jagged1 portions, Jagged1^EGF8-11^ and Jagged1^CRD^ (Figures S7a-h), show both samples do not change their oligomeric state at different concentrations (Table 2; Figures S7b and S7f). While Jagged1^EGF8-11^ is flexible (Figure S7d), Jagged1^CRD^ is compact and globular (Figure S7h). The measured *D*_max_ of 240 Å indicates monomeric Jagged1^fe,HA^ is partially backfolded, as a fully elongated Jagged1 ectodomain would have a maximum dimension of 585 Å (see Methods). In agreement with the SAXS data, the XL-MS analysis suggest that the extracellular region of Jagged1 is back-folded instead of fully extended (Figures 2a and 2b). The detected distance restraints arise from either intra- or intermolecular Jagged1^fe^ interactions, as Jagged1^fe^ may be dimerizing in this experiment. To isolate the intramolecular cross-links from the ambiguous intra- and intermolecular cross-links we repeated the cross-linking experiment with Jagged1^fe,HA^ and separated monomeric Jagged1^fe,HA^ from cross-linked Jagged1^fe,HA^ oligomers by SEC (Figure S2b) and analyzed the cross-links of both fractions by MS. The data indicate that four regions of the Jagged1 extracellular segment (C2-EGF2, EGF5-6, EFG9-12 and CRD) are in proximity within the same Jagged1^fe,HA^ molecule, as most identified cross-links are present in the monomeric (as well as in the oligomeric) fraction (Figures 4g and S2c). Most of these intramolecular cross-links are also found in the Notch1fe-Jagged1^fe,*wt*^ and Notch1^fe^-Jagged1^fe,HA^ XL-MS datasets, indicating that these intramolecular cross-links are independent of Notch1^fe^ binding.

We used SPR to verify that the regions identified by XL-MS interact directly. Constructs consisting of the Jagged1 regions C2-EGF3, EGF8-11 and CRD reveal direct interactions between Jagged1^C2-EGF3^ and Jagged1^EGF8-11^, and between Jagged1^C2-EGF3^ and Jagged1^CRD^, supporting the XL-MS results. The interactions are weak as covalent dimerization by Fc-fusion was required to measure binding. Fc-Jagged1^EGF8-11^ and Fc-Jagged1^CRD^ bound to Jagged1^C2-EGF3,*wt*^ with a *K*_D,app_ of 0.34 µM and 0.93 µM, respectively (Figures 4h, 4i, S1k and S1l). The C2-EGF3 region is required and sufficient for these interactions. Both Fc-Jagged1^EGF8-11^ and Fc-Jagged1^CRD^ do not interact with Jagged1^EGF5-CRD^ that is lacking the C2-EGF3 region (Figures 4h and 4i) and affinities are similar for larger constructs that include the C2-EFG3 region, *i*.*e*. Jagged1^C2-EGF7^, Jagged1^C2-EGF13^ and Jagged1^fe^ (Table 1). In addition, the Jagged1 high-affinity mutations (Luca *et al*., 2017) do not affect this interaction (Table 1). Taken together, the SPR and XL-MS data indicate that the EGF8-11 and CRD regions interact intramolecularly with the C2-EGF3 region within the Jagged1 molecule.

**Figure 5.**
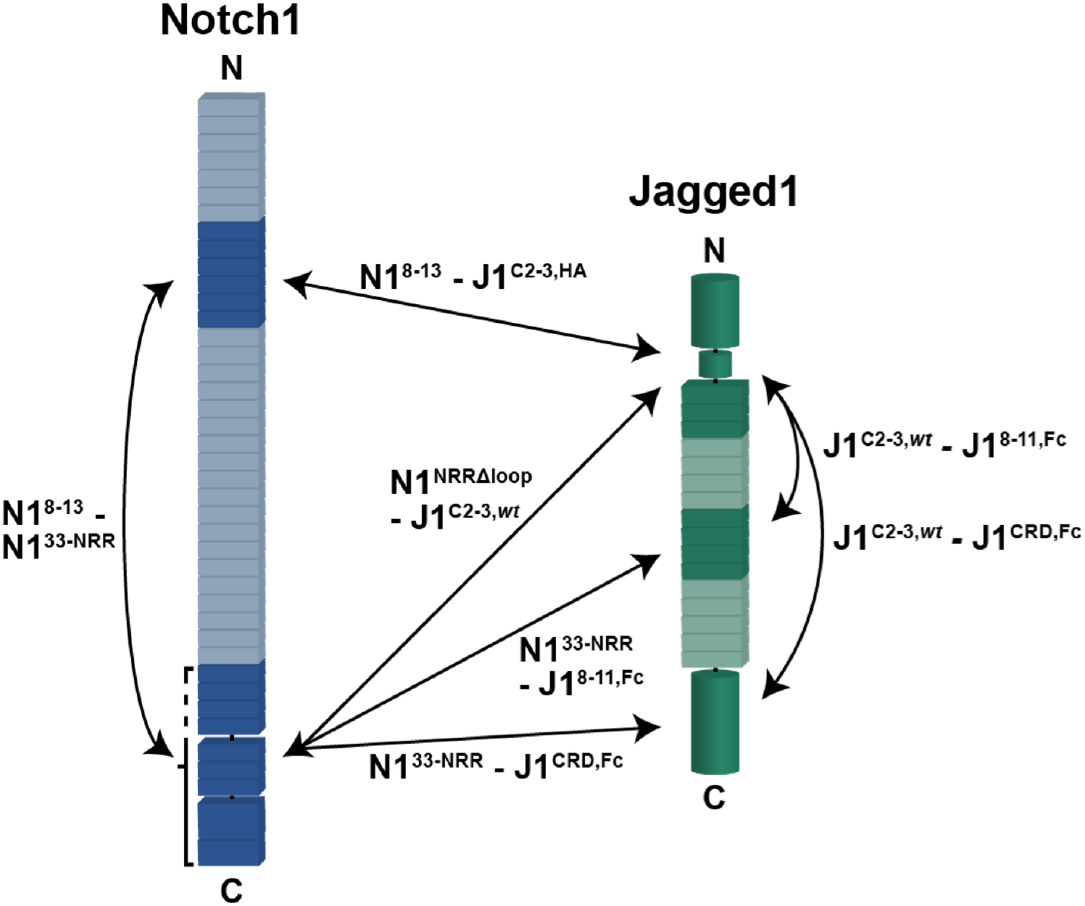
Summary of the uncovered and validated direct interactions. Inter- and intra-molecular interactions based on the XL-MS and quantitative-interaction experiments are indicated by double arrows.

## Discussion

Two regions in Notch, EGF11-12 and NRR, have been widely studied due to their critical role in Notch signaling (Brou *et al*., 2000; Handford *et al*., 2018; Logeat *et al*., 1998; Lovendahl *et al*., 2018; Mumm *et al*., 2000; Rebay *et al*., 1991) and represent the minimal requirements for ligand-dependent Notch activation (Cordle *et al*., 2008b; Gordon *et al*., 2007). Transcellular ligand binding at the Notch1 EGF8-12 site, positioned far away from the NRR in the primary sequence, and subsequent Notch1-ligand endocytosis generate a mechanical pulling force (Chowdhury *et al*., 2016; Gordon *et al*., 2015; Lovendahl *et al*., 2018; Luca *et al*., 2017; Meloty-Kapella *et al*., 2012; Seo *et al*., 2016; Wang and Ha, 2013) that could be transmitted via EGF13-36 to the NRR where it triggers a conformational change to expose the S2 site to proteolytic cleavage (Brou *et al*., 2000; Gordon *et al*., 2007; Mumm *et al*., 2000). Ligand binding in cis can inhibit Notch activation (Del Á lamo *et al*., 2011; D’Souza *et al*., 2008; Sprinzak *et al*., 2010), while it was recently shown that it could also stimulate Notch activation (Nandagopal *et al*., 2019), although it is not clear if and how endocytosis plays a direct role in this setting. These studies raise the question of how the different regions within Notch1 and Jagged1 interact.

Here we show that the Jagged1 C2-EGF3 segment is in close proximity to the Notch1 NRR in the Notch1^fe^-Jagged1^fe^ complex, that Notch1^EGF8-13^ and Notch1^NRR^ can interact directly with the C2-EGF3 region in Jagged1, and that Notch1^EGF8-13^ interacts with Notch1^EGF33-NRR^. We confirm that the Notch1 ectodomain has regions of flexibility (Pei and Baker, 2008; Sharma *et al*., 2013; Weisshuhn *et al*., 2016), which suggests that the EGF8-13 and the NRR segments in Notch1 can interact intramolecularly. The binding of Jagged1 C2-EGF3 to the membrane proximal Notch1 NRR fits well with the previously shown lipid-binding role of the Jagged1 C2 domain and the requirement of C2-lipid binding for optimal Notch activation (Chillakuri *et al*., 2013; Suckling *et al*., 2017). The various segments have different interaction strengths. The interaction of the Notch1 ectodomain and that of Jagged1 is weak but strengthened by a pulling force (Luca *et al*., 2017). The mutation of five residues in the Jagged1 C2 domain increases the affinity of the Jagged1 ectodomain for the Notch1 ectodomain to 1 µM (Figure 2d), indicating that the Jagged1 C2 domain plays an important role in the interaction with Notch1. Surprisingly, the measured interaction between Notch1^NRRΔloop^ and Jagged1^C2-EGF3^ also has a *K*_D_ of about 1 µM and is not dependent on the high-affinity mutations (Figure 2e). While this interaction may be influenced in the SPR experiment by an avidity effect, arising from dimerization of the NRR, the interaction measured between Notch1^NRR^ and Jagged1^C2-EGF3,HA^ in solution using MST also shows a *K*_D_ of around 1 µ]M (Figure 2f). The interaction of the larger Notch1^EGF33-NRR^ with Jagged1^C2-EGF3^ shows a similar affinity with a *K*_D_ of 0.5 µM measured by MST (Figure 2f), whereas it is 30-fold weaker in the surface-based SPR method (Figure S4b and Table 1), which indicates that the context of this interaction may be important. Taken together, these data show that the NRR in the Notch1 ectodomain is in direct contact to the Jagged1 C2-EGF3 region in the Notch1^fe^-Jagged1^fe^ complex and suggest that ligand binding is directly coupled to Notch activation or regulation.

The setting at the cell surface or between two cells may dictate how Notch1 and Jagged1 interact. In our experiments we cannot discriminate between cis and trans interactions, and it may be possible we see both types of interactions simultaneously. For example, the interaction of the membrane proximal regions, *i*.*e*. Notch1 EGF33-NRR and Jagged1 CRD, seems more likely in a cis setting with both molecules expressed on the same cell. At the same time, the receptor and the ligand may undergo homomeric interactions on the cell surface which influences Notch signaling further (Hicks *et al*., 2002; Kelly *et al*., 2010; Sakamoto *et al*., 2005; Shimizu, 2002; Vooijs *et al*., 2004; Xu *et al*., 2015). Besides the C2-EGF3 region, we have identified additional Jagged1 segments, namely EGF8-11 and CRD, that interact intermolecularly with Notch1 EGF33-NRR as well as intramolecularly with Jagged1 C2-EGF3, and these regions could have a role in the clustering of Jagged1 and the Notch1-Jagged1 complex on, or between, cells. It is currently not clear whether the Notch1 NRR - Jagged1 C2-EGF3 and Notch1 EGF8-13 - NRR interactions are common features for the Notch and DSL family members. Interestingly, despite differences in domain composition, these three regions are present in all members, *i*.*e*. all Notch paralogs contain the EGF8-13 and NRR segments and all DSL ligands have the C2-EGF3 region in common. Our data indicate that a mosaic of interaction sites is present, both on Notch1 and on Jagged1. Targeting these interactions may reveal their role in Notch signaling and could have potential for therapeutic applications to treat Notch-associated disorders.

## Methods

### Generation of constructs and mutagenesis

Notch1 and Jagged1 constructs were generated by polymerase chain reaction (PCR) using mouse Notch1 (Addgene 41728), human Notch1 (kind gift of Dr. Warren Pear, Univ. of Pennsylvania) and mouse Jagged1 (Image clone 6834418) as templates. All constructs are mouse version unless stated otherwise. Notch1^fe^ (residue numbers 19-1717) was subcloned in pUPE106.03 (U-Protein Express BV, cystatin secretion signal peptide, N-terminal His6-tag), Notch1^fe^ (19-1728, human version), Notch1^EGF8-13^ (294-526), Notch1^EGF22-27^ (828-1058), Notch1^EGF33-36^ (1267-1426), Notch1^EGF33-NRR^ (1267-1717), Notch1 ^NRR^ (1446-1717) with and without its unstructured loop (1622-1659), Jagged1^fe^ (31-1067), Jagged1^C2-EGF3^ (31-334), Jagged1^C2-EGF7^ (31-485), Jagged1^C2-EGF13^ (31-741), Jagged1^EGF5-13^ (374-741), Jagged1 ^EGF5-CRD^ (374-1067), Jagged1^EGF8-11^ (487-665), Jagged1^CRD^ (857-1067) were subcloned in pUPE107.03 (U-Protein Express BV, cystatin secretion signal peptide, C-terminal His6-tag). Jagged1 mutations (S32L, R68G, D72N, T87R, Q182R) based on (Luca *et al*., 2017) were introduced using Q5 Site-Directed Mutagenesis to generate Jagged1^fe,HA^ (31-1067) and Jagged1^C2-EGF3,HA^ (31-334) constructs. In several figures, Notch1 and Jagged1 constructs are referred to as N1 and J1, respectively, and EGF repeats are referred to as their number, *i*.*e*. J1C2-3 for Jagged1^C2-EGF3^.

### Large-scale expression and purification

Constructs were transiently expressed in N-acetylglucoaminyltransferase I-deficient (GnTI-) Epstein-Barr virus nuclear antigen 1 (EBNA1)-expressing HEK293 cells growing in suspension (U-Protein Express BV). The medium was harvested six days after transfection, cells were spun down by 10 minutes of centrifugation at 1000x g, and cellular debris was spun down for 15 minutes at 4000x g. For human Notch1^fe^ used in the SEC-MALS experiment, the supernatant was concentrated fivefold and diafiltrated against 25 mM 4-(2-hydroxyethyl)-1-piperazineethanesulfonic acid (HEPES) pH 8.0, 500 mM NaCl and 2 mM CaCl2 (IMAC A) using a Quixstand benchtop system (GE Healthcare) with a 10 kDa molecular weight cutoff (MWCO) membrane. Cellular debris were spun down for 10 min at 9500x g and the concentrate was filtered with a glass fiber prefilter (Minisart, Sartorius). Protein was purified by Nickel-nitrilotriacetic acid (Ni-NTA) affinity chromatography and eluted with a mixture of 60% IMAC A and 40% of 25 mM HEPES pH 8.0, 500 mM NaCl, 500 mM imidazole, 2 mM CaCl_2_ (IMAC B). For all other constructs and experiments, cells were spun down by 10 minutes of centrifugation at 1000x g, cellular debris was spun down for 15 minutes at 4000x g, and protein was directly purified by Ni Sepharose excel (GE Healthcare) affinity chromatography. Protein was eluted with a mixture of 60% of IMAC C (same as IMAC A, except pH 7.4) and 40% of IMAC D (same as IMAC B, except pH 7.4), or with 100% of IMAC D. SEC was performed on either a Superose6 10/300 increase (GE Healthcare) or a Superdex200 10/300 increase (GE Healthcare) equilibrated in SEC buffer (20 mM HEPES pH 7.4, 150 mM NaCl, 2 mM CaCl2). Protein purity was evaluated by sodium dodecyl sulphate-polyacrylamide gel electrophoresis (SDS-PAGE) and Coomassie staining. Protein was concentrated and then stored at −80 °C.

### Protein Cross-linking with PhoX

XL-MS was performed according to a previously optimized protocol (Klykov *et al*., 2018). The optimal cross-linker concentration was established with SDS-PAGE. Cross-linking reactions were performed in triplicates with equimolar inputs of each protein for Notch1^fe^-Jagged1^fe,*wt*^ and for Notch1^fe^- Jagged1^fe,HA^. Purified proteins at concentration of 10 µM were incubated together in cross-linking buffer (20 mM HEPES pH 7.8, 150 mM NaCl and 1.5 mM MgCl2) for 10 minutes followed by adding of PhoX (Thermo) to a final concentration of 1 mM. The sample mixtures were filtered through MWCO 10 kDa filters (Vivaspin) into 10 mM Tris pH 7.5 in a 3:1 ratio (*v:v*) to a final volume of 25 µl. Prior to protein digestion, samples were deglycosylated overnight with Deglycosylation Mix II (NEBB). After deglycosylation, urea was added to a final concentration of 8 M followed by addition of Tris(2-carboxyethyl)phosphine (TCEP) and chloroacetamide to a final concentration of 10 mM and 40 mM respectively. Samples were incubated at 37°C for 1 hour and then proteolytic digestion was performed with LysC (Wako) for 4 hours and trypsin (Promega) overnight. Resulting peptide mixtures were desalted with Oasis HLB plates (Waters), dried and stored at −80°C until further use.

### Automated Fe(III)-IMAC-Based Enrichment

Cross-linked peptides were enriched with Fe(III)-NTA 5 µL in an automated fashion using the AssayMAP Bravo Platform (Agilent Technologies). Fe(III)-NTA cartridges were primed with 250 µL of 0.1% TFA in ACN and equilibrated with 250 µL of loading buffer (80% ACN/0.1% TFA). Samples were dissolved in 200 µL of loading buffer and loaded onto the cartridge. The columns were washed with 250 µL of loading buffer, and the cross-linked peptides were eluted with 25 µL of 10% ammonia directly into 25 µL of 10% formic acid. Samples were dried down and stored in 4 °C until subjected to LC-MS. For LC-MS analysis the samples were resuspended in 10% formic acid.

### Liquid Chromatography Mass Spectrometry and data analysis

All mass spectrometry data was acquired using an UHPLC 1290 system (Agilent Technologies) coupled on-line to an Orbitrap Fusion Lumos mass spectrometer (Thermo Scientific). Peptides were trapped (Dr. Maisch Reprosil C_18_, 3 µm, 2 cm x 100 µm) prior to separation on an analytical column (Agilent Poroshell EC-C_18_, 2.7 µm, 50 cm x 75 µm). Trapping was performed by flushing in buffer A (0.1% *v:v* formic acid in water) for 10 min. Reversed phase separation was performed across a gradient of 10 % to 40 % buffer B (0.1% *v:v* formic acid in 80% *v:v* ACN) over 90 min at a flow-rate of approximately 300 nL/min. The instrument was operated in data-dependent MS2 mode with MS1 spectra recorded in the range 350-1400 Th and acquired in the Orbitrap at a resolution of 60,000 with an AGC of 4 × 10^5^ and a maximum injection time of 50 ms. For MS2, the cycle time was set to 3 s with charge state inclusion set to 3-8 for the enriched fraction and 2-8 for the flow-through. Dynamic exclusion was set to 12 s at 1.4 Th mass deviation. Stepped HCD was performed with the Ion Trap at NCE = 35 (+/− 10%) and acquired in the Orbitrap at a resolution of 30,000 with AGC set at 1 × 10^5^ maximum injection time to 120 ms. The cross-linked peptides were analyzed with Thermo Proteome Discoverer (2.3.0.522) with incorporated XlinkX/PD nodes (Klykov *et al*., 2018). The analysis was run with standard parameters in NonCleavable mode at 1 % False Discovery rate (FDR) at the level of the CSM and Cross-link tables against a manually created database with the target proteins and 200 random decoy entries. As fixed modification Carbamidomethyl (C) was set and as variable modification Oxidation (M), Acetyl (protein N-term), and Asn→Asp (N) (H_-1_ N_-1_ O). As cross-linking reagent PhoX (C_8_ H_3_ O_5_ P) was set. Only cross-links detected in 2 out of 3 replicates were used for further analysis. The normal and mono-linked peptides were analyzed with MaxQuant (1.6.17.0) (Cox and Mann, 2008). The analysis was run with standard settings applied using the same database to search the spectra. As fixed modification Carbamidomethyl (C) was set and as variable modification Oxidation (M), Acetyl (protein N-term), PhoX Tris (K) (C_12_ H_14_ N O_8_ P), PhoX H2O (K) (C_8_ H_5_ O_6_ P) and Asn *→* Asp (N) (H_-1_ N_-1_ O). Further downstream analysis and visual representation of the results was performed with the R scripting and statistical environment (Ihaka and Gentleman, 1996) using Circos (Krzywinski *et al*., 2009) for data visualization. The mass spectrometry raw data, result/search files and the annotated spectra have been deposited to the ProteomeXchange Consortium via the PRIDE partner repository (Perez-Riverol *et al*., 2019) with the dataset identifier PXD023072.

### Integrative modeling and docking of Notch1 NRR and Jagged1 C2-EGF3

To the crystal structure of Notch1 NRR described here (PDB: 7ABV), the missing flexible loop modelled with trRosetta (Yang *et al*., 2020) was added, *i*.*e*. residues 1622-1659. A structure of mouse Jagged1 C2-EGF3 was generated by homology modelling in ITASSER (Yang *et al*., 2015) based on the rat high-affinity Jagged1 variant template (PDB: 5UK5; Luca et al., 2017). Next, Notch1 NRR with the added loop and Jagged1 C2-EGF3 were docked together with three XL-MS based restraints from these regions and defined as 5-25 Å distance restraints in the HADDOCK2.4 webserver (Figure S5; van Zundert *et al*., 2016). The loop was defined as fully flexible and the resulting outputs of the complex were examined in terms of scores with the emphasis on the biological relevance and restraints energy violations. UCSF ChimeraX (Pettersen *et al*., 2020) was used for visualization.

### Surface plasmon resonance

SPR ligand constructs subcloned in-frame in pUPE107.62 (cystatin secretion signal peptide, C-terminal biotin acceptor peptide-tag followed by a C-terminal His6-tag) were biotinylated in HEK293 cells by co-transfection with E. coli BirA biotin ligase with a sub-optimal secretion signal (in a pUPE5.02 vector), using a DNA ratio of 9:1 (sample:BirA, m/m). Additional sterile biotin (100 µL of 1 mg/mL HEPES-buffered biotin per 4 mL HEK293 culture) was supplemented to the medium. Protein was purified from the medium by Ni Sepharose excel (GE Healthcare) affinity chromatography. Purity was evaluated by SDS-PAGE and Coomassie staining. C-terminally biotinylated proteins were spotted on a P-STREP SensEye (Ssens) chip with a Continuous Flow Microspotter (CFM, Wasatch Microfluidics) using an 8×6 format. SEC buffer with 0.005% Tween-20 was used as a spotting buffer and the coupling was quenched using 1 mM biotin in SEC buffer. Proteins were therefore C-terminally coupled to the chip to ensure a native topology. Surface plasmon resonance experiments were performed on an MX96 SPRi instrument (IBIS Technologies). Analytes in SEC buffer were flowed over the sensor chip, and SEC buffer with 0.005% Tween-20 was used as a running buffer. Temperature was kept constant at 25 °C. The data was analyzed using SprintX (IBIS Technologies) and Prism (Graphpad) and modeled with a 1:1 Langmuir binding model to calculate the *K*_D_ and the maximum analyte binding (B_max_). Since the NRR dimerizes, and bound with positive cooperativity to Jagged1^C2-EGF3^ when it was used as an analyte, we fitted SPR equilibrium binding plots using a Hill equation with a Hill coefficient of 2. For the experiments in which full regeneration could not be achieved, the subsequent analyte injections were not zeroed in order to keep the B_max_ constant (see Figures S1d, S1e, S1i and S1l).

### Microscale Thermophoresis

Jagged1^C2-EGF3,HA^ in SEC buffer was labelled with NT-547 dye (NanoTemper Technologies) according to the manufacturer’s instructions. Unlabelled Notch1^EGF33-NRR^ and Notch1^NRR^ in SEC buffer were serially diluted from 50 µM to 3.0 nM (Notch1^EGF33-NRR^) or 1.5 nM (Notch1^NRR^) and incubated with 50 nM labelled Jagged1^C2-EGF3,HA^ in the presence of 0.025% Tween-20 for 15 minutes at room temperature. Samples were transferred to Standard Treated Capillaries (NanoTemper Technologies) and run at 50 % excitation power on a Monolith NT.115 (NanoTemper Technologies) at a constant temperature of 25 °C. *K*_D_ was determined according to the law of mass action using the program MO Affinity Analysis (NanoTemper Technologies) and results were plotted using Prism (Graphpad).

### Small-angle X-ray scattering

Notch1^fe^ SEC-SAXS experiments were carried out at the European Synchrotron Radiation Facility (ESRF) beamline BM29. 500 µL of 8.1 µM human Notch1^fe^ were loaded on a Superose6 10/300 increase column (GE Healthcare) equilibrated in SEC buffer, via a high-performance liquid chromatography system (Shimadzu). A stable background signal was confirmed before measurement. Measurements were performed at room temperature at a flow rate of 0.5 mL/min. SAXS data was collected at a wavelength of 0.99 Å using a sample-to-detector (Pilatus 1M, Dectris) distance of 2.85 m. The scattering of pure water was used to calibrate the intensity to absolute units. 2000 frames of 2 s each were collected and data reduction was performed automatically using the EDNA pipeline (Incardona et al., 2009). Frames with a stable *R*_g_ (+/− 10 %) and buffer frames were selected for further analysis using Chromixs (Panjkovich and Svergun, 2018). Data was analyzed in Primus (Konarev *et al*., 2003) and Scatter (Förster *et al*., 2010), and results were plotted in Prism (Graphpad). The maximum dimension of 1027 Å for a theoretical elongated Notch1 ectodomain was calculated as follows: an average of 27 Å for the 36 EGF repeats (Luca *et al*., 2017) and 55 Å for the NRR (Gordon *et al*., 2009a).

Jagged1^fe,HA^ SEC-SAXS experiments were carried out at the Diamond Light Source (DLS) beamline B21 operating at an energy of 12.4 keV and using a sample-to-detector (Eigen 4M, Dectris) distance of 4.01 m. 45 µL of 42 µM Jagged1^fe,HA^ were loaded on a Superose6 3.2/300 increase (GE Healthcare) equilibrated in SEC buffer, via a HPLC system (Agilent). A stable background signal was confirmed before measurement. Measurements were performed at room temperature at a flow rate of 0.075 mL/min. The scattering of pure water was used to calibrate the intensity to absolute units. 620 frames of 3 s each were collected and data reduction was performed automatically using the DAWN pipeline (Filik *et al*., 2017). Frames with a stable *R*_g_ and buffer frames were selected for further analysis using Chromixs (Panjkovich and Svergun, 2018). Data was analyzed in Primus (Konarev *et al*., 2003) and Scatter (Förster *et al*., 2010), and results were plotted in Prism (Graphpad).

Jagged1^EGF8-11^, Jagged1^CRD^ and Jagged1^fe^ batch SAXS experiments were carried out the DLS beamline B21 operating at an energy of 12.4 keV and using a sample-to-detector (Eigen 4M, Dectris) distance of 4.01 m. The scattering of pure water was used to calibrate the intensity to absolute units. Data reduction was performed automatically using the DAWN pipeline (Filik *et al*., 2017). Frames were averaged after being manually inspected for radiation damage, the scattering of the SEC buffer was subtracted, and intensities were normalized by the concentration. Data was analyzed in Primus (Konarev *et al*., 2003) and Scatter (Förster *et al*., 2010), and results were plotted in Prism (Graphpad). The maximum dimension of 585 Å for a theoretical elongated Jagged1 ectodomain was calculated as follows: 160 Å for the C2-EGF3 region as measured from its crystal structures (Chillakuri *et al*., 2013; Luca *et al*., 2017), an average of 27 Å for each of the remaining 13 EGF domains (Luca *et al*., 2017), and 75 Å as determined for the C-terminal CRD by SAXS (Figure S7G).

### Multi-Angle Light Scattering

SEC-MALS was performed using a Superose6 10/300 increase (GE Healthcare) column for Notch1^fe^ (human version) or a Superdex 10/300 increase (GE Healthcare) column for Jagged1^fe,HA^ and Notch1^NRR^, equilibrated in SEC buffer. For molecular weight characterization, light scattering was measured with a miniDAWN TREOS multi-angle light scattering detector (Wyatt Technology) connected to a RID-10A differential refractive index monitor (Shimadzu) for quantitation of the protein concentration. Chromatograms were collected, analyzed and processed on the ASTRA software suite (Wyatt Technology). A dn/dc of 0.1800 was calculated for Notch1^fe^ based on 6 N-glycosylation sites of the oligo-mannose type and 55 O-glycosylation sites (2 sugar moieties per site), 0.1814 for Jagged1^fe,HA^ based on 9 N-glycosylation sites and 16 O-glycosylation sites (4 O-glucosylation sites extended with 2 xylose moieties each, and 12 O-fucosylation sites), and 0.1828 for Notch1^NRR^ based on 2 N-glycosylation sites.

### Crystallization and data collection

The Notch1 NRR was crystallized by sitting-drop vapour diffusion at 18 °C, by mixing 200 nL of protein solution containing a mixture of Notch1^NRR^ and Jagged1 ^C2-EGF3,HA^ at 8.5 mg/mL in SEC buffer, and 100 nL of reservoir solution, composed of 2.0 M sodium chloride and 0.1 M sodium acetate pH 4.6. The protein solution was deglycosylated beforehand using EndoHf 1:100 (*v:v*) overnight at room temperature in SEC buffer. The crystal was harvested and flash-cooled in liquid nitrogen in the presence of reservoir solution supplemented with 25% glycerol. The dataset was collected at 100 K at the DLS beamline I03 (*λ* = 1.06998 Å).

### Structure solution and refinement

The data was processed by the autoPROC pipeline (Vonrhein *et al*., 2011) consisting of XDS (Kabsch, 2010), POINTLESS (Evans, 2006), AIMLESS (Evans and Murshudov, 2013), CCP4 (Winn *et al*., 2011) and STARANISO (Tickle *et al*., 2018). The structure was solved by molecular replacement by searching for one copy of PDB ID 3ETO (Gordon *et al*., 2009a). After molecular replacement, the model was improved by manual model building in Coot (Emsley *et al*., 2010) and refinement with REFMAC (Murshudov *et al*., 2011). Validation was performed using MolProbity (Chen *et al*., 2010).

### Analytical ultracentrifugation

SV-AUC experiments were carried out in a Beckman Coulter Proteomelab XL-I analytical ultra-centrifuge with An-60 Ti rotor (Beckman) at 40,000 revolutions per minute (r.p.m.). Jagged1^fe,HA^ at 5 µM and at 20 µM were measured in SEC buffer at 20 °C. Either 12 mm (5 µM sample) or 3 mm (20 µM sample) centerpieces with quartz windows were used. Absorbance was determined at 280 nm using SEC buffer as a reference. A total of 800 scans per cell were collected and analyzed in continuous c(s) mode in SEDFIT (Schuck, 2000). Buffer density and viscosity were determined with SEDNTERP as 1.0061 g/mL and 0.010314 Pa·s, respectively.

## Acknowledgments

We thank the staff of the DLS beamline B21, the staff of the ESRF beamline BM29, Dimphna H. Meijer and Mercedes Ramírez-Escudero for assistance with SAXS data collection, and the staff of the DLS beamline I03 for assistance with X-ray diffraction data collection. We thank Nadia Leloup for help with Notch1 SEC-MALS experiments and Dominique Thies-Weesie for help with SV-AUC experiments. This project has received funding from: the European Research Council (ERC) under the European Union’s Horizon 2020 research and innovation programme with grant agreement No. 677500 (to B.J.C.J.); the research programme TA with project number 741.018.201 (to R.A.S.), which is partly financed by the Dutch Research Council (NWO); and the European Union Horizon 2020 programme INFRAIA project Epic-XS project 823839 (to R.A.S.). This work benefited from access to the Amsterdam NKI, an Instruct-ERIC centre, with financial support provided by Instruct-ERIC (PID 10025). This work has been supported by iNEXT (PID 6764), funded by the Horizon 2020 programme of the European Union.

## Data and materials availability

The mass spectrometry raw data, result/search files and the annotated spectra have been deposited to the ProteomeXchange Consortium via the PRIDE partner repository with the dataset identifier PXD023072. All SAXS data is made available at the Small Angle Scattering Biological Data Bank (SASBDB) with the accession codes SASDJG8 (Monomeric Notch1^fe^), SASDJ38 (Monomeric Jagged1^fe,HA^), SASDJ48 (5.3 µM Jagged1^fe,HA^), SASDJ58 (11 µM Jagged1^fe,HA^), SASDJ68 (21 µM Jagged1^fe,HA^), SASDJ78 (42 µM Jagged1^fe,HA^), SASDJ88 (29 µM Jagged1^EGF8-11^), SASDJ98 (58 µM Jagged1^EGF8-11^), SASDJA8 (115 µM Jagged1^EGF8-11^), SASDJB8 (230 µM Jagged1^EGF8-11^), SASDJC8 (21 µM Jagged1^CRD^), SAS-DJD8 (42 µM Jagged1^CRD^), SASDJE8 (83 µM Jagged1^CRD^), SASDJF8 (167 µM Jagged1^CRD^). Coordinates and structure factors for S1-cleaved mouse Notch1 NRR have been deposited to the Protein Data Bank (PDB) with accession code 7ABV.

## Contributions

M.R.Z. and B.J.C.J. designed the experiments and interpreted all data; M.R.Z., J.P.M., M.J.K. and A.G. generated constructs and purified proteins; O.K. performed the MS experiments and data analysis; M.R.Z., J.P.M. and M.J.K performed the SPR experiments; M.R.Z. and M.J.K. performed the MST and crystallization experiments; M.R.Z. and J.P.M. performed the SAXS experiments; M.R.Z. performed the X-ray diffraction, SEC-MALS, and SV-AUC experiments; B.J.C.J. and R.A.S. supervised the project; M.R.Z., O.K., R.A.S. and B.J.C.J. wrote the manuscript with input from all authors; B.J.C.J. conceived the project.

