## Supplementary figures for "Notch-Jagged signaling complex defined by an interaction mosaic"

**This PDF file includes:**

Figures S1 to S7
Table S1

**Other supplementary materials for this manuscript include the following:**

Datasets S1 to S2

### Sensorgrams and MST traces related to Fig. 2

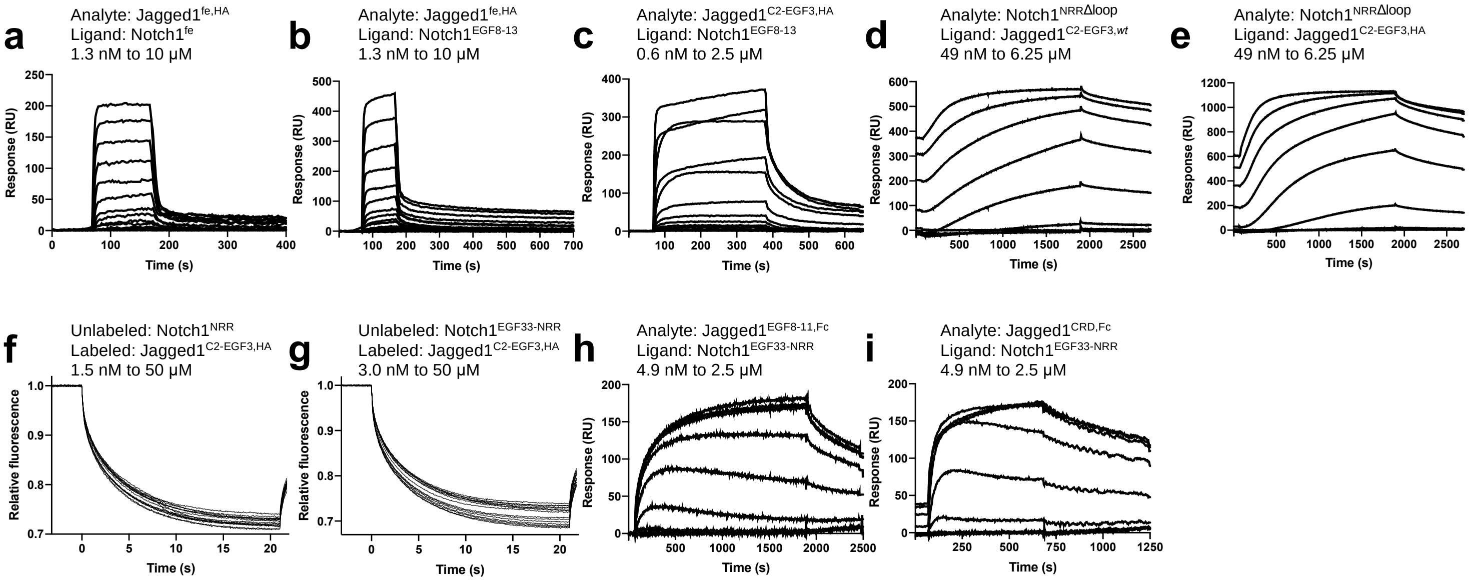


### Sensorgram related to Fig. 3

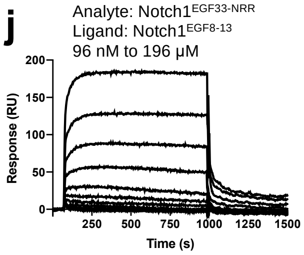


### Sensorgrams related to Fig. 4

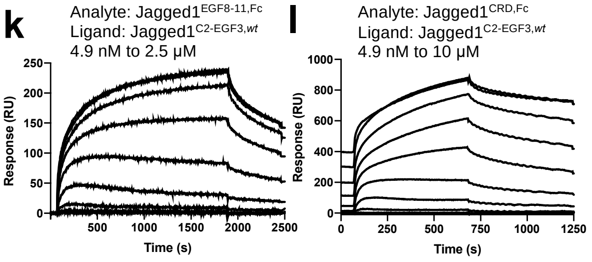


**Fig. S1. Sensorgrams and MST traces, related to Fig. 2-4. a-i** Sensorgrams and MST traces related to **Fig. 2**, with sensorgrams of Jagged1$^{fe,HA}$ binding **a** to Notch1$^{fe}$ and **b** to Notch1$^{EGF8\text{-}13}$, **c** Jagged1$^{C2\text{-}EGF3,HA}$ binding to Notch1$^{EGF8\text{-}13}$, Notch1$^{NRR\Delta loop}$ binding **d** to Jagged1$^{C2\text{-}EGF3,\mathit{wt}}$ and **e** to Jagged1$^{C2\text{-}EGF3,HA}$, MST traces of **f** Notch1$^{NRR}$ binding to Jagged1$^{C2\text{-}EGF3,HA}$ and **g** Notch1$^{EGF33\text{-}NRR}$ binding to Jagged1$^{C2\text{-}EGF3,HA}$, sensorgrams of **h** Jagged1$^{EGF8\text{-}11,Fc}$ binding to Notch1$^{EGF33\text{-}NRR}$ and **i** Jagged1$^{CRD,Fc}$ binding to Notch1$^{EGF33\text{-}NRR}$. **j** Sensorgram related to Fig. 3, with Notch1$^{EGF33\text{-}NRR}$ binding to Notch1$^{EGF8\text{-}13}$. **k,l** Sensorgrams related to Fig. 4, with **k** Jagged1$^{EGF8\text{-}11,Fc}$ binding to

Jagged1$^{C2\text{-}EGF3,wt}$ and **I** Jagged1$^{CRD,Fc}$ binding to Jagged1$^{C2\text{-}EGF3,wt}$. The concentration range used in the experiment is indicated in all panels.

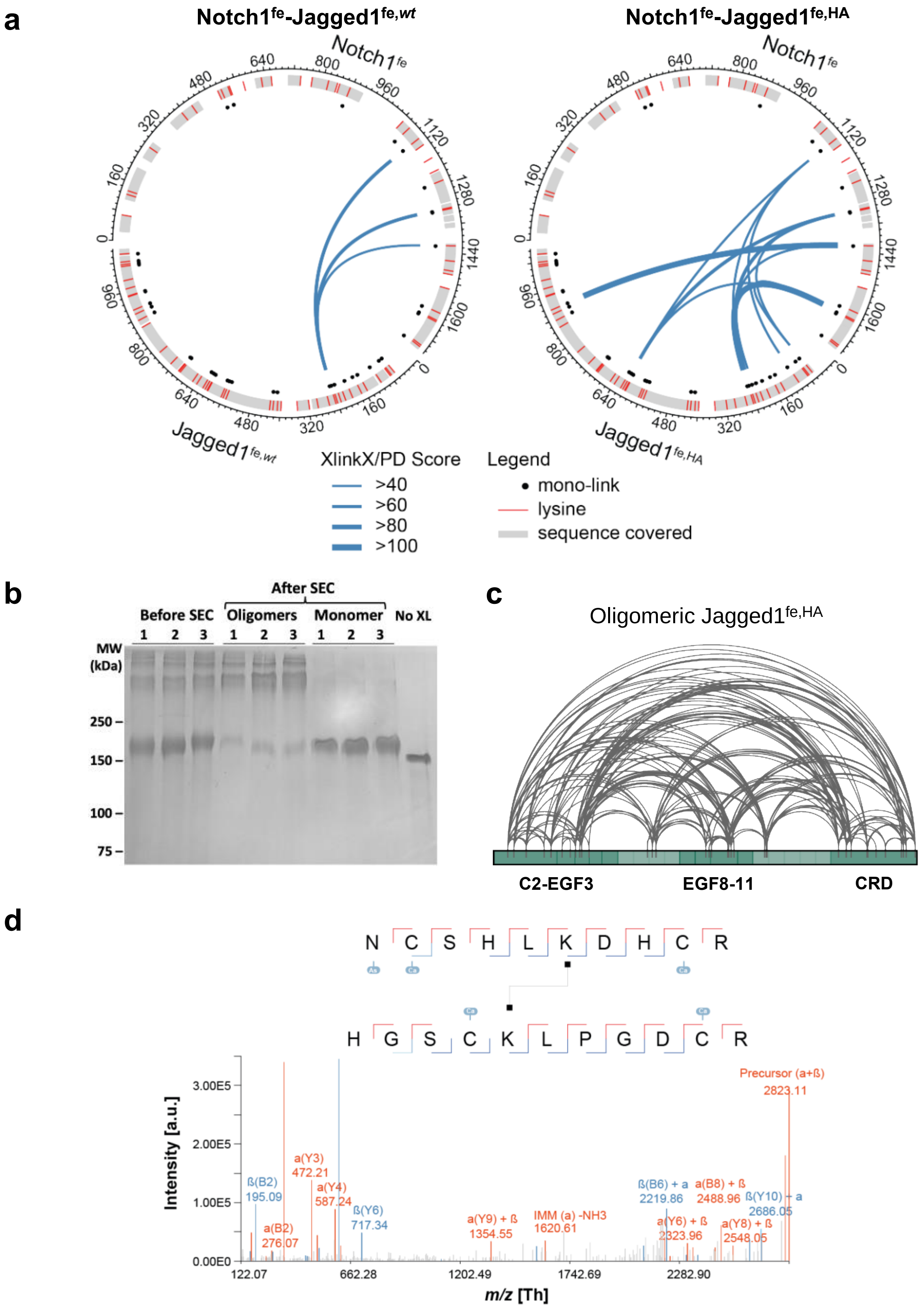
a
Notch1fe-Jagged1fe,wt
Notch1fe-Jagged1fe,HA
Notch1fe
Jagged1fe,wt
Jagged1fe,HA
XlinkX/PD Score
>40
>60
>80
>100
Legend
mono-link
lysine
sequence covered
b
After SEC
Before SEC
Oligomers
Monomer
No XL
MW (kDa)
250
150
100
75
c
Oligomeric Jagged1fe,HA
C2-EGF3
EGF8-11
CRD
d
N C S H L K D H C R
H G S C K L P G D C R
Intensity [a.u.]
m/z [Th]
Precursor (a+ß) 2823.11
ß(B2) 195.09
a(B2) 276.07
a(Y3) 472.21
a(Y4) 587.24
ß(Y6) 717.34
a(Y9) + ß 1354.55
IMM (a) -NH3 1620.61
ß(B6) + a 2219.86
a(Y6) + ß 2323.96
a(B8) + ß 2488.96
a(Y8) + ß 2548.05
ß(Y10) + a 2686.05
3.00E5
2.00E5
1.00E5
0.00E0
122.07
662.28
1202.49
1742.69
2282.90

**Fig. S2. Additional information related to cross-linking mass-spectrometry experiments, related to Fig. 2,4. a** Circular plots indicating the inter-links, by XlinkX/Proteome Discoverer score, and mono-links identified in the cross-linking experiment of Notch1$^{fe}$-Jagged1$^{fe,\textit{wt}}$ (left) and Notch1$^{fe}$-Jagged1$^{fe,HA}$ (right). The sequence covered in the peptide identification is indicated. **b** Coomassie-stained SDS-PAGE showing the cross-linked oligomeric and monomeric Jagged1 fractions purified by size exclusion chromatography in triplicate. The monomer fractions are well separated from the oligomeric fraction. **c** Overview of the detected distance constraints from the XL-MS experiments for oligomeric Jagged1$^{fe,HA}$. The detected distance constraints for monomeric Jagged1$^{fe,HA}$ are shown in **Fig. 4g**. **d** Example mass spectrum of an identified cross-link.

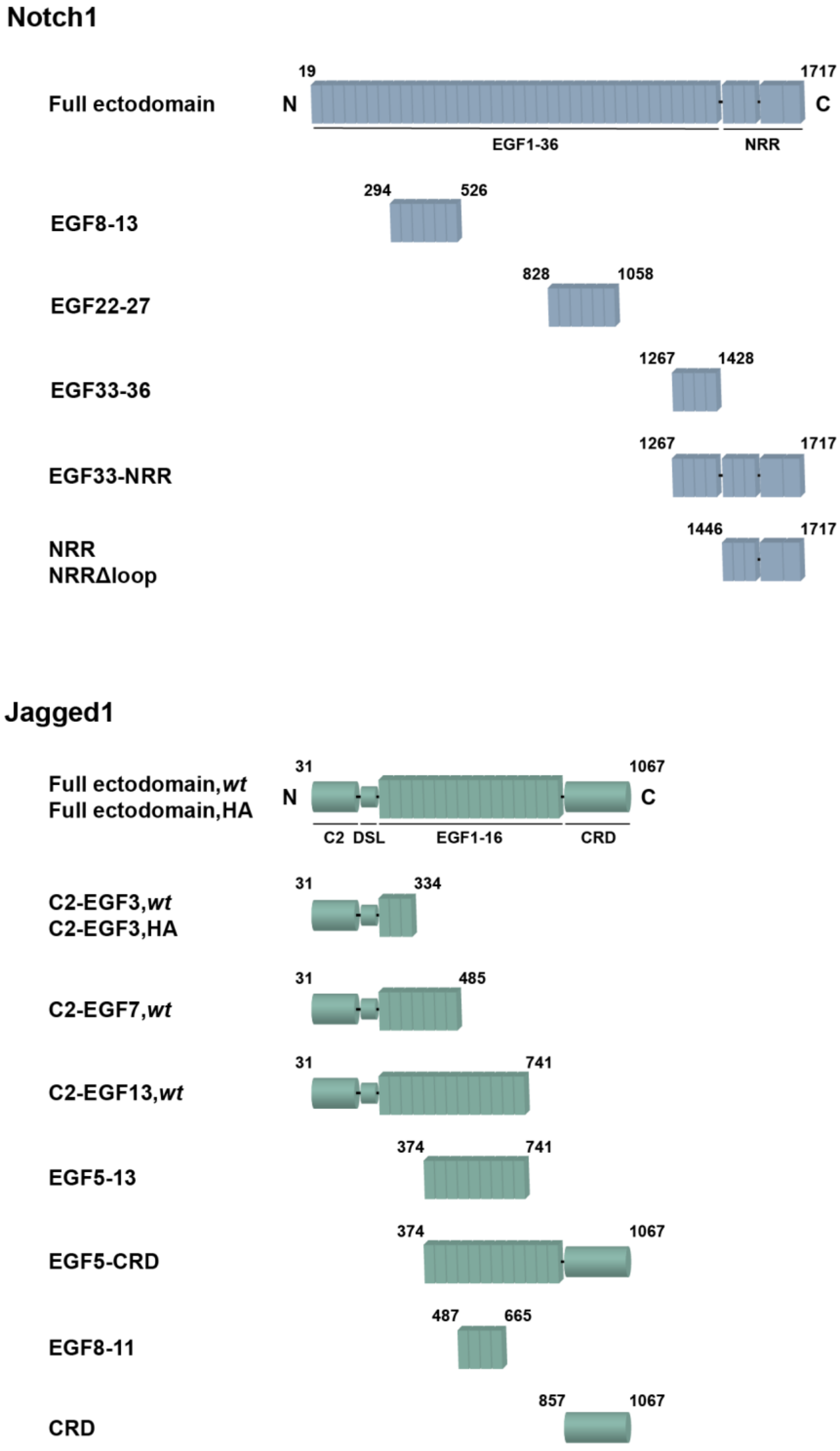


**Fig. S3. Domain composition and main constructs generated, related to Fig. 2-4.**

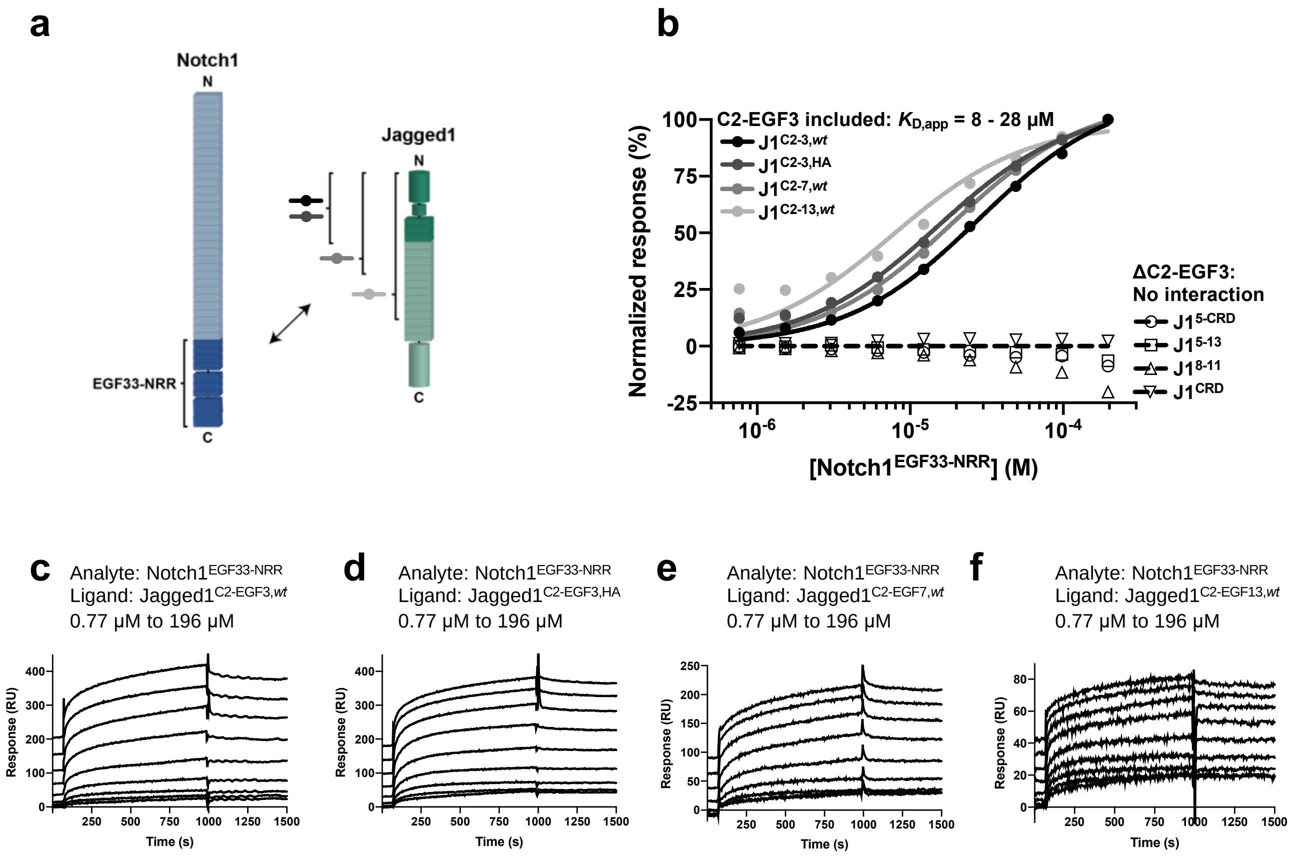


**Fig. S4. The C2-EGF3 domain of Jagged1 is necessary and sufficient for Notch1 EGF33-NRR interaction, related to Fig. 2. a** Schematic representation of the interactions reported in panels **(b-f)**. **b** SPR equilibrium binding plots of Notch1$^{EGF33\text{-}NRR}$ to Jagged1$^{C2\text{-}EGF3,wt}$ (black), Jagged1$^{C2\text{-}EGF3,HA}$ (dark grey), Jagged1$^{C2\text{-}EGF7,wt}$ (grey), Jagged1$^{C2\text{-}EGF13,wt}$ (light grey), Jagged1$^{EGF5\text{-}CRD}$ (open circle), Jagged1$^{EGF5\text{-}13}$ (open square), Jagged1$^{EGF8\text{-}11}$ (open triangle) and Jagged1$^{CRD}$ (open inverted triangle). **c-f** Corresponding SPR sensorgrams, with Notch1$^{EGF33\text{-}NRR}$ binding **c** to Jagged1$^{C2\text{-}EGF3,wt}$, **d** to Jagged1$^{C2\text{-}EGF3,HA}$, **e** to Jagged1$^{C2\text{-}EGF7,wt}$, and **f** to Jagged1$^{C2\text{-}EGF13,wt}$.

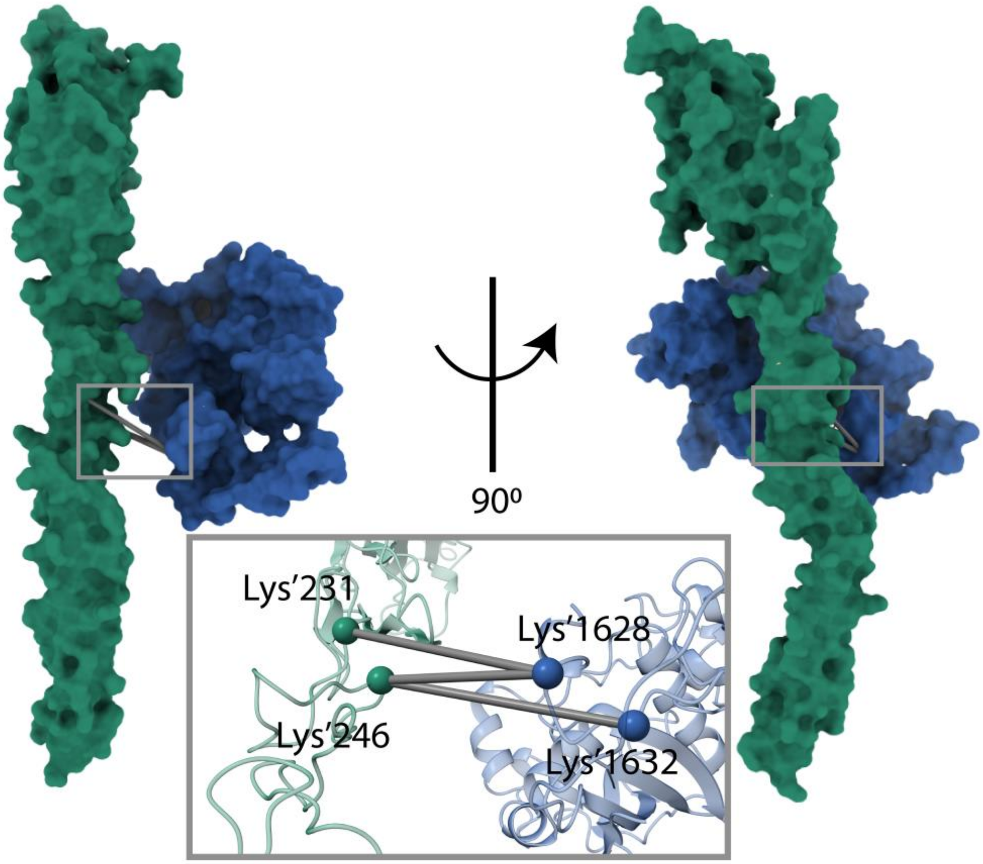


**Fig. S5. Modelling of Notch1 NRR - Jagged1 C2-EGF3 complex, related to Fig. 2.** Docking of the Notch1 NRR - Jagged1 C2-EGF3 complex using the structure of Notch1 NRR described here (blue) and that of Jagged1 C2-EGF3 (green; PDB: 5UK5) and based on cross-links obtained by XL-MS. The two structures are slightly separated from each other to indicate the cross-links.

**a**

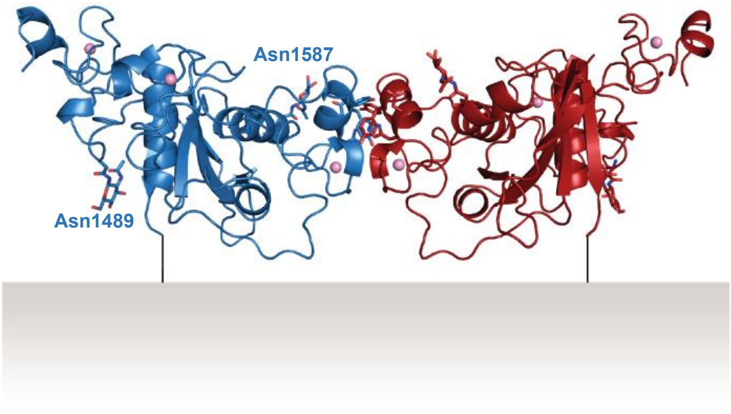


**b**

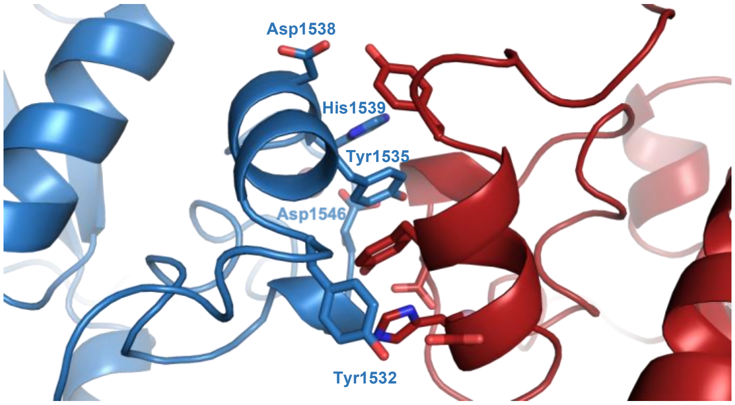


**c**

| | S1-cleaved mouse Notch1 NRR |
|---|---|
| **Data collection** | |
| Space group | $P\ 4_2\ 2_1\ 2$ |
| Cell dimensions | |
| *a*, *b*, *c* (Å) | 65.72, 65.72, 161.68 |
| *α*, *β*, *γ* (°) | 90, 90, 90 |
| Resolution (Å) | 60.89 - 2.06 (2.30 - 2.06) |
| No. observed reflections | 195430 (8934) |
| No. unique reflections | 15144 (757) |
| $R_{merge}$ | 0.105 (1.431) |
| Mean I/σI | 14.6 (1.7) |
| $CC_{1/2}$ | 0.999 (0.770) |
| Completeness (spherical, %) | 66.9 (12.6) |
| Completeness (ellipsoidal, %) | 91.5 (64.6) |
| Ellipsoidal resolution limits (Å) [direction] | 2.44 [a*]<br>2.44 [b*]<br>2.06 [c*] |
| Redundancy | 12.9 (11.8) |
| **Refinement** | |
| Resolution (Å) | 60.89 - 2.06 |
| $R_{work}/R_{free}$ (%) | 19.53 / 23.39 |
| Average *B*-factors (Å$^2$) | |
| Protein | 57.7 |
| Glycans/ions | 94.5 |
| Waters | 46.6 |
| R.M.S. deviations | |
| Bond lengths (Å) | 0.0084 |
| Bond angles (°) | 1.52 |
| Ramachandran (%) | |
| Favored | 95.61 |
| Allowed | 4.39 |
| Outliers | 0 |
| Molprobity score | 2.06 |

**Fig. S6. Structure of the S1-cleaved mouse Notch1 NRR, related to Fig. 3. a** Proposed orientation of the Notch1 NRR dimer with respect to the cell surface. **b** Key residues at the dimerization interface are indicated. **c** Data collection and refinement statistics. Highest resolution shell in parentheses.

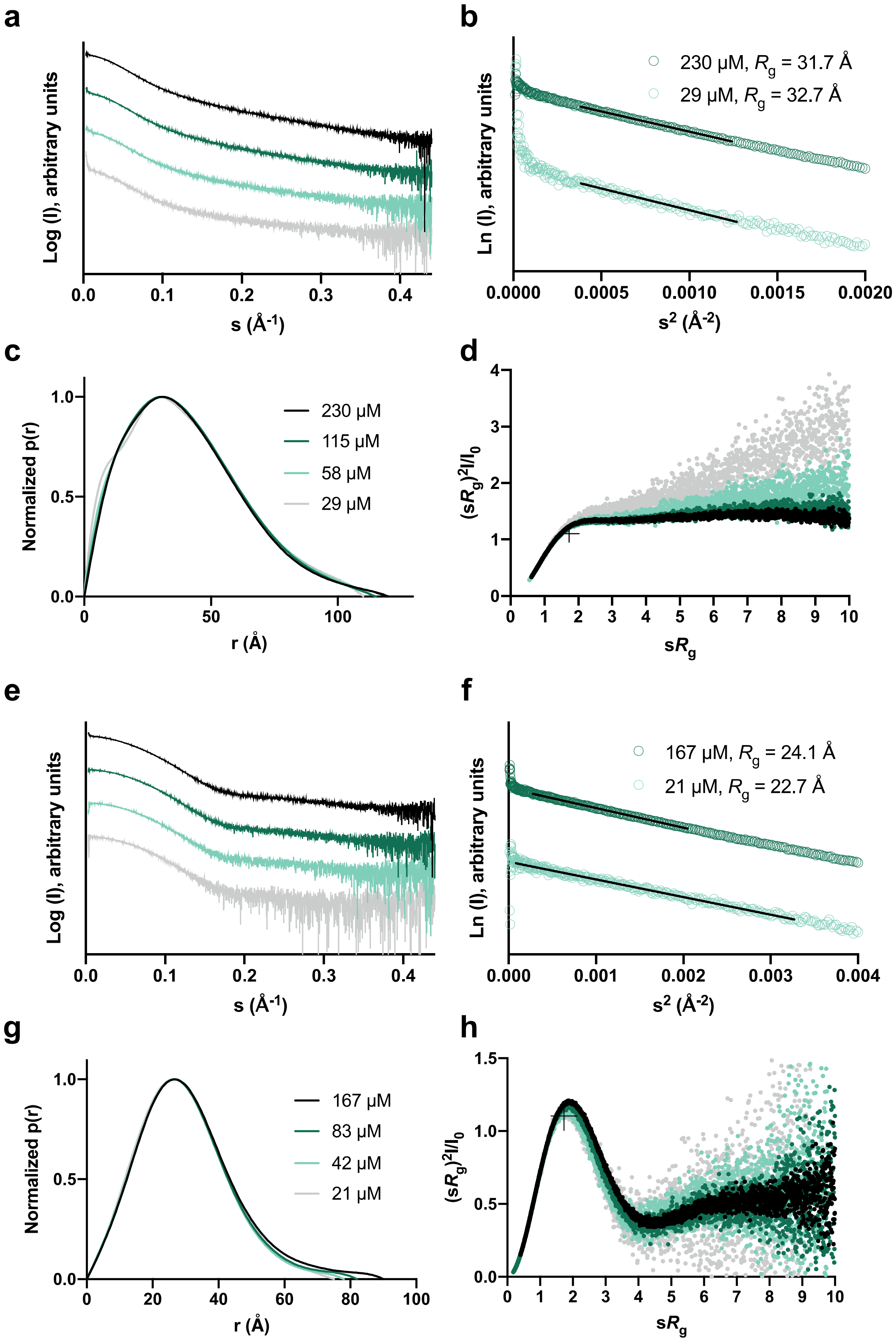
a
Log (I), arbitrary units
0.0 0.1 0.2 0.3 0.4
s (Å-1)
b
230 µM, Rg = 31.7 Å
29 µM, Rg = 32.7 Å
Ln (I), arbitrary units
0.0000 0.0005 0.0010 0.0015 0.0020
s2 (Å-2)
c
Normalized p(r)
1.0 0.5 0.0
230 µM
115 µM
58 µM
29 µM
0 50 100
r (Å)
d
(sRg)2I/I0
4 3 2 1 0
0 1 2 3 4 5 6 7 8 9 10
sRg
e
Log (I), arbitrary units
0.0 0.1 0.2 0.3 0.4
s (Å-1)
f
167 µM, Rg = 24.1 Å
21 µM, Rg = 22.7 Å
Ln (I), arbitrary units
0.000 0.001 0.002 0.003 0.004
s2 (Å-2)
g
Normalized p(r)
1.0 0.5 0.0
167 µM
83 µM
42 µM
21 µM
0 20 40 60 80 100
r (Å)
h
(sRg)2I/I0
1.5 1.0 0.5 0.0
0 1 2 3 4 5 6 7 8 9 10
sRg

**Fig. S7. Jagged1$^{EGF8-11}$ and Jagged1$^{CRD}$ have distinct structural properties, related to Fig. 4.** **a-d** Structural analysis of Jagged1$^{EGF8-11}$ from batch SAXS, including a Log (I) versus s plot, **b** Guinier plot with black lines indicating the fits used to derive the $R_g$, **c** pair distance distribution function and **d** dimensionless Kratky plot with crosshairs indicating the peak position for a globular protein. **e-h** Structural analysis of Jagged1$^{CRD}$ from batch SAXS, including **e** Log (I) versus s plot, **f** Guinier plot with black lines indicating the fits used to derive the $R_g$, **g** pair distance distribution function and **h** dimensionless Kratky plot with crosshairs indicating the peak position for a globular protein.

**Table S1. Description of the files uploaded to the PRIDE repository, related to Fig. 2.**

| Filename | Description |
|---|---|
| 20190316_L1_Ag6_Klyko001_SA_NOTCH_PhoX_MT1.raw<br>20190316_L1_Ag6_Klyko001_SA_NOTCH_PhoX_MT2.raw<br>20190316_L1_Ag6_Klyko001_SA_NOTCH_PhoX_MT3.raw | Triplicate measurements of the crosslinked and PhoX enriched fraction for the mouse Notch1 - mouse high-affinity mutant Jagged1 full ectodomain complex. |
| 20190316_L1_Ag6_Klyko001_SA_NOTCH_PhoX_MT_Asn-_AspSites.txt<br>20190316_L1_Ag6_Klyko001_SA_NOTCH_PhoX_MT_evidence.txt<br>20190316_L1_Ag6_Klyko001_SA_NOTCH_PhoX_MT_peptides.txt<br>20190316_L1_Ag6_Klyko001_SA_NOTCH_PhoX_MT_PhoX H2OSites.txt<br>20190316_L1_Ag6_Klyko001_SA_NOTCH_PhoX_MT_PhoX TrisSites.txt<br>20190316_L1_Ag6_Klyko001_SA_NOTCH_PhoX_MT.pdf | MaxQuant output tables and annotated spectra. |
| 20190316_L1_Ag6_Klyko001_SA_NOTCH_PhoX_MT_Crosslinks.txt<br>20190316_L1_Ag6_Klyko001_SA_NOTCH_PhoX_MT_CSMs.txt<br>20190316_L1_Ag6_Klyko001_SA_NOTCH_PhoX_MT_CSMs.pdf | Proteome Discoverer XlinkX/PD output tables and annotated spectra. |
| 20190316_L1_Ag6_Klyko001_SA_NOTCH_PhoX_WT1.raw<br>20190316_L1_Ag6_Klyko001_SA_NOTCH_PhoX_WT2.raw<br>20190316_L1_Ag6_Klyko001_SA_NOTCH_PhoX_WT3.raw | Triplicate measurements of the crosslinked and PhoX enriched fraction for the mouse Notch1 - mouse wild-type Jagged1 full ectodomain complex. |
| 20190316_L1_Ag6_Klyko001_SA_NOTCH_PhoX_WT_Asn-_AspSites.txt<br>20190316_L1_Ag6_Klyko001_SA_NOTCH_PhoX_WT_evidence.txt<br>20190316_L1_Ag6_Klyko001_SA_NOTCH_PhoX_WT_peptides.txt<br>20190316_L1_Ag6_Klyko001_SA_NOTCH_PhoX_WT_PhoX H2OSites.txt<br>20190316_L1_Ag6_Klyko001_SA_NOTCH_PhoX_WT_PhoX TrisSites.txt<br>20190316_L1_Ag6_Klyko001_SA_NOTCH_PhoX_WT.pdf | MaxQuant output tables and annotated spectra. |
| 20190316_L1_Ag6_Klyko001_SA_NOTCH_PhoX_WT_Crosslinks.txt<br>20190316_L1_Ag6_Klyko001_SA_NOTCH_PhoX_WT_CSMs.txt<br>20190316_L1_Ag6_Klyko001_SA_NOTCH_PhoX_WT_CSMs.pdf | Proteome Discoverer XlinkX/PD output tables and annotated spectra. |
| 20190923_L1_Ag1_Klyko001_SA_JagNotch_PhoX_FT_MT1.raw<br>20190923_L1_Ag1_Klyko001_SA_JagNotch_PhoX_FT_MT2.raw<br>20190923_L1_Ag1_Klyko001_SA_JagNotch_PhoX_FT_MT3.raw | Triplicate measurements of the flow-through (i.e. not cross-linked & mono-linked) fraction for the mouse Notch1 - mouse high-affinity mutant Jagged1 full ectodomain complex. |
| 20190923_L1_Ag1_Klyko001_SA_JagNotch_PhoX_FT_MT_Asn-_AspSites.txt<br>20190923_L1_Ag1_Klyko001_SA_JagNotch_PhoX_FT_MT_evidence.txt<br>20190923_L1_Ag1_Klyko001_SA_JagNotch_PhoX_FT_MT_peptides.txt<br>20190923_L1_Ag1_Klyko001_SA_JagNotch_PhoX_FT_MT_PhoX H2OSites.txt<br>20190923_L1_Ag1_Klyko001_SA_JagNotch_PhoX_FT_MT_PhoX TrisSites.txt<br>20190923_L1_Ag1_Klyko001_SA_JagNotch_PhoX_FT_MT.pdf | MaxQuant output tables and annotated spectra. |
| 20190923_L1_Ag1_Klyko001_SA_JagNotch_PhoX_FT_WT1.raw<br>20190923_L1_Ag1_Klyko001_SA_JagNotch_PhoX_FT_WT2.raw<br>20190923_L1_Ag1_Klyko001_SA_JagNotch_PhoX_FT_WT3.raw | Triplicate measurements of the flow-through (i.e. not cross-linked & mono-linked) fraction for the mouse Notch1 - mouse wild-type Jagged1 full ectodomain complex. |
| 20190923_L1_Ag1_Klyko001_SA_JagNotch_PhoX_FT_WT_Asn-_AspSites.txt<br>20190923_L1_Ag1_Klyko001_SA_JagNotch_PhoX_FT_WT_evidence.txt<br>20190923_L1_Ag1_Klyko001_SA_JagNotch_PhoX_FT_WT_peptides.txt<br>20190923_L1_Ag1_Klyko001_SA_JagNotch_PhoX_FT_WT_PhoX H2OSites.txt<br>20190923_L1_Ag1_Klyko001_SA_JagNotch_PhoX_FT_WT_PhoX TrisSites.txt<br>20190923_L1_Ag1_Klyko001_SA_JagNotch_PhoX_FT_WT.pdf | MaxQuant output tables and annotated spectra. |

**Dataset S1 (separate file). This annotated excel file contains the Crosslinks table of XlinkX/PD, broken up in inter- and intra-links, related to Fig. 2.** The original Crosslinks and CSM tables can be found in the PRIDE repository.

**Dataset S2 (separate file). This annotated excel file contains the Site specific tables of MaxQuant for the PhoX:Tris and PhoX;H2O monolinks, related to Fig. 2.** The original tables plus the Evidence and Peptide tables can be found in the PRIDE repository.
